# PRER: A Patient Representation with Pairwise Relative Expression of Proteins on Biological Networks

**DOI:** 10.1101/2020.06.16.153999

**Authors:** Halil İbrahim Kuru, Mustafa Buyukozkan, Oznur Tastan

## Abstract

Changes in protein and gene expression levels are often used as features to predictive models such as survival prediction. A common strategy to aggregate information on individual proteins is to integrate the expression information with biological networks. We propose a novel patient representation in this work where we integrate proteins’ expression levels with the protein-protein interaction (PPI) networks. Patient representation with PRER (<u>P</u>airwise <u>R</u>elative <u>E</u>xpressions with <u>R</u>andom walks) uses the neighborhood of a protein to capture the dysregulation patterns in protein abundance. Specifically, PRER computes a feature vector for a patient by comparing the source protein’s protein expression level with other proteins’ levels in its neighborhood. This neighborhood of the source protein is derived using a biased random-walk strategy on the network. We test PRER’s performance through a survival prediction task in 10 different cancers using random forest survival models. PRER representation yields a statistically significant predictive performance in 9 out of 10 cancer types when compared to a representation based on individual protein expression. We also identify important proteins that are not important in the models trained with the expression values but emerge as predictive in models trained with PRER features. The set of identified relations provides a valuable collection of biomarkers with high prognostic value. PRER representation can be used for other complex diseases and prediction tasks that use molecular expression profiles as input. PRER is freely available at: https://github.com/hikuru/PRER

## 1 Introduction

With the advances in sequencing technologies, large-scale molecular profiling of patients has become possible. The comprehensive profiling of cancer patients, along with the available patient clinical data, presents an opportunity to gain deeper insights into cancer and develop prediction tools for diagnostic, prognostic, and therapeutic purposes. Machine learning has been an instrumental tool for realizing this aim. In these studies, patients are often represented with their molecular data, such as gene expression profiles encoded as numerical feature vectors. For example, Yuan et al. [1] assess the utility of different types of molecular changes for survival prediction. While using miRNA, protein, or mRNA expression, they use these entities’ expression values as input. Others follow a similar approach for different clinical outcome prediction tasks [2, 3, 4].

Genes and proteins interact to carry out their functional roles in the cell, and phenotypes arise from these functional interactions. Based on this basic principle, alternative approaches where the patient molecular profiles are integrated with the prior knowledge of molecular interactions have been proposed (reviewed in [5] and [6]). Taking into account the network of interactions help to aggregate the signals attached to each protein or gene in a biologically principled way. Integration of the expression levels of genes/proteins and their interactions are used in multiple studies [7, 8, 9, 10, 11, 11]. Chuang et al. [8] was among the first. They identify discriminant and highly altered subnetworks of interactions using gene expression data and use the activity summaries of genes on these subnetworks as features for metastasis prediction. By assessing the association of pathways and transcription factors with overall survival as opposed to individual genes, Crijns et al. [10] identify signaling pathways and transcription factors that contribute to the clinical outcome of ovarian cancer. Taylor et al. [9] integrate a PPI network with a co-expression profile and report that the genes with dysregulated neighbors in that PPI network are potential prognostic markers. NetBank [12] uses gene expressions and prior knowledge network to rank genes according to their relevance to the outcome of pancreatic cancer. These studies aggregate dysregulations in a subnetwork, pathway, or network by summing or diffusing them in the network without the relative expression changes. Different from the above methods, Wang and Liu [13] et al. use the topological importance of the proteins in the network to reweight them in random survival forest sampling. There are also methods that have used this idea for other types of omic profiles [5]. For example, Hofree et al. [7] integrates mutation data with PPI network for patient stratification. They employ the network propagation idea to diffuse the mutation information along with a PPI and use that representation for patient stratification.

A limited number of studies use the pairwise comparisons of molecular measurements instead of the aggregation of expression levels. Geman et al. [14] report a method that uses the pairwise ranks of mRNA expression levels for classifying gene expression profiles in tumor identification, disease detection, and treatment response. Magen et al. [15] use pairwise combinations of expression dysregulations to predict survival-related gene pairs. These methods, however, do not make use of the prior knowledge available in the biological networks.

In this work, we explore a method that combines the two ideas discussed above; network integration and pairwise comparison of expression levels. Pairwise Rank Expressions with Random walks (PRER) is a novel molecular representation method that considers the relative expression of a protein within its neighborhood on the PPI network. A given protein’s neighborhood is defined based on a biased random walk search on the PPI network. PRER also allows interpretability. The pairwise relationships of interacting neighborhood molecules offer a direct interpretation of molecular dysregulation patterns in the context of known protein interactions. We also present methods to analyze pairs that are predictive because of their pairwise comparisons.

We computed PRER representation using protein expression data obtained from patient tumors for survival prediction in ten cancers provided by the Cancer Genome Atlas (TCGA) project [16]. When compared to the representation of patients with their protein expression features, PRER yields a statistically significant improvement in 9 of the 10 cancer types. PRER also outperform better against two additional competitive methods. Additionally, PRER unveils predictive features concerning the known PPIs. We also investigate proteins that are deemed significant solely based on their interactions.

## 2 Methods

### 2.1 PRER Feature Representation

PRER constructs a vector-based patient representation for subsequent prediction tasks by integrating the patients’ molecular expression profiles and the PPI network. The molecular expressions can be the mRNA expressions or protein expressions. Since not all the transcript expression changes are reflected as changes at the protein expression level, in this work, we choose to use protein expression data as input to PRER.

Let *G* = (*V, E*) be the given PPI network, where *V* is the set of vertices representing the proteins, and *E* is the set of edges that exist between proteins if known to interact. Let *U* ⊂ *V* be the proteins for which protein expression values are available for all patients in the data set. The nodes with the protein expression data, *U*, constitute the source proteins, and we will denote the number of such proteins with *m*. Given *G*, *U*, and patient expression data over *U*, the output of PRER for a patient *k* is a feature vector, *x*^(*k*)^ ∈ *R^s^*, that contains the pairwise comparisons encoded with 1 and −1’s. Here, *s* denotes the size of the pairwise comparisons, which will be clarified in the following sections. Below we detail the steps of PRER.

#### Step 1. Obtaining a Protein’s Neighborhood on the Protein Interaction Network

For each source protein in *U*, we first define a neighborhood, *N*_*u*_, which is the set of proteins proximal to the source protein *u* on *G*. To obtain the neighborhood of a node in the graph, a set of random walks is generated. For every source node *u* ∈ *U*, we sample neighbors of the source node with a strategy similar to the one in the node2vec [17] algorithm. A random walk with a fixed length of *l* starting at source node *u* is generated based on the following distribution:

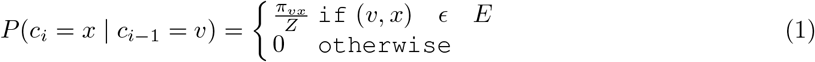

Here, *c*_*i*_ denotes *i*_*th*_ node in the walk and *c*_0_ = *u*. **Z** is the normalization constant. *P* (*c*_*i*_ = *x c*_*i*−1_ = *v*) is the transition probability on edge (*v, x*), where the current node is *v*, the next node to visit is *x*, and the previous node is *t*. The transition probability depends on the function *π*, and it is defined as:

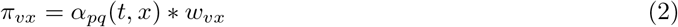

where *w*_*vx*_ is the edge weight between nodes *v* and *x*. However, in this work, we use an unweighted PPI network and, thus, we set *w*_*vx*_ = 1. *α*_*pq*_(*t, x*) is the random walk bias which is defined by equation 3 based on the parameters *p* and *q* and the shortest path distance between nodes *t* and *x*, *d*_*tx*_

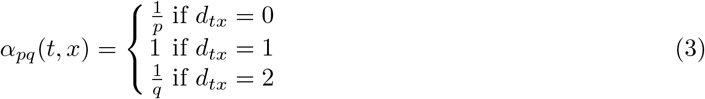

This bias controls the different search strategies to sample the next visited nodes. We use two different search methods: depth-first sampling (DFS) and breadth-first sampling (BFS), as in [17]. BFS samples the nodes from the nearby nodes, whereas DFS samples the nodes sequentially by gradually increasing the distance from a source node. *p* and *q* parameters control the connection between BFS and DFS approaches. With a high *q* value, sampled nodes in the random walk are aligned to BFS and get a local view over the source node. A small *q* value aligns random walk to DFS to explore a global view of the network. *p* controls the chance of revisiting the nodes. A high value of *p* decreases the probability of sampling the already visited nodes, while a small value of *p* aligns random walk to return the source node.

This biased random walk strategy has two additional parameters: (i) walk length *l* and (ii) the number of random walks *r*. We select these parameters based on the parameter sensitivity analysis at node2vec [17]. The parameters *p* and *q* are used as *p* = 0.25*, q* = 0.25 in our random walk generation. When *p* = 1*, q* = 1 uniform random walks are generated without any bias as stated in Grover and Leskovec. A small *q* value is used to bias the random walks to capture the network’s global view, while a small *p* value is used to capture the community around the source node *u*. With the given values, random walks are inclined to see the communities inside the network. By using fixed-length (*l* = 100) random walks, we sample a neighborhood for a given source node, *u*. To be consistent and to decrease the variance, multiple random walks per source node are applied so that different neighborhoods are sampled for each node. We sampled random walks 18 times, and these are stored in *W*_*B*_ (see Figure 1). The frequency of nodes in the multiple neighborhoods is calculated, and the nodes involved in more than one random walk are selected as the neighborhood genes.

**Fig. 1:**
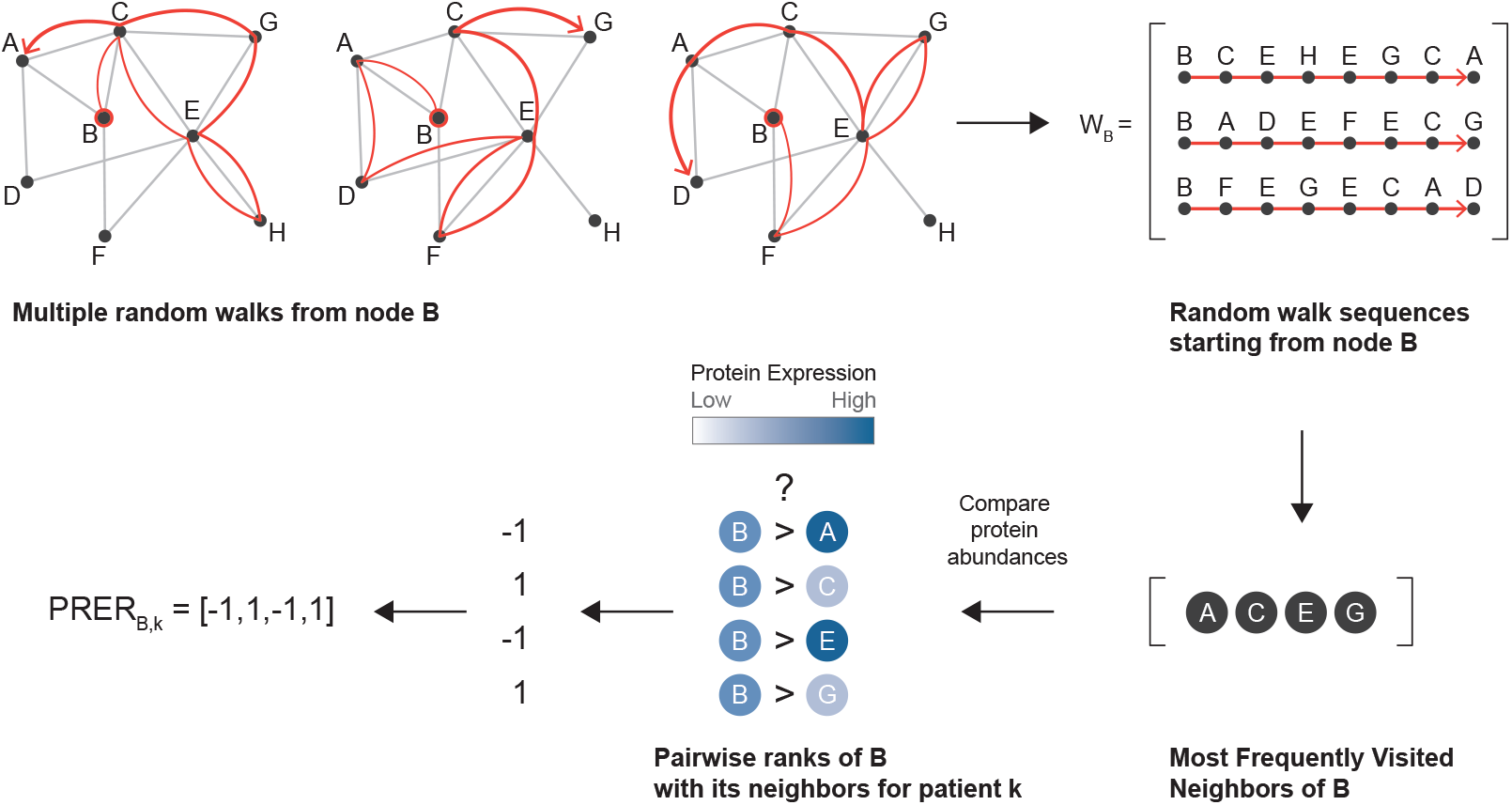
Illustration to show how the PRER representation is obtained for a single source node, node B. The nodes in the graph are proteins, edges exist if they interact in the PPI network. First, several random walks are generated that starts at node B as in [17]. These random walks are stored in *W*_*B*_ and used to define the neighborhood of *B*, *N*_*B*_. Only the most frequently visited nodes are included in the set of neighbors of B. Then, the pairwise comparison of the neighborhood proteins in terms of their protein expression quantities is used to form a representation of the patient for node B and its neighborhood. The figure shows the features generated for a single protein. This procedure is repeated for all source proteins, and the resulting vectors are concatenated.

#### Step 2. Feature Representation based on Pairwise Rank of Neighborhood Genes

At the end of step one, we arrive at the neighborhood of the protein *i*, which we denote as *N*_*i*_. Some neighbors lack measurements, and we define the subset of neighbor proteins with accompanying measurements as *M*_*i*_ = ∈ *N*_*i*_ ⋂ *U*. Next, for a protein *i*, we generate pairwise rank features with every protein *i* ∈ *M*_*i*_ as follows.

Let 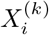 and 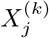 denote the expression quantities for protein *i* and *j* for patient *k*. Protein *i* is the source protein, and protein *j* is a protein in the neighborhood of *i*. The pairwise rank expression representations (PRER) for this patient is defined as:

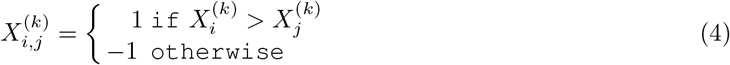

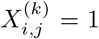 indicates that the molecule *i* is more upregulated with respect to molecule *j* for this patient, whereas 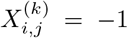 indicates otherwise. For every *i* in *U* and every *j* in *M*_*i*_, we define a pairwise rank order for the protein pair. If the protein *i*’s phosphorylated state or states are measured, their comparison with *i* is also included. Since the current PPIs do not account for the phosphorylated state (e.g., STATPY705) when we create features for the phosphorylated state of the protein, we use the neighbors of the unphosphorylated (e.g., STAT3) node in the PPI network.

This representation constitutes a nonlinear interaction feature mapping among original features that aims to capture expression dysregulations among interacting proteins. This representation is a nonparametric rank statistics. The rank-based statistics are widely used to obtain a robust statistical analysis. For example, Kendall tau is a robust correlation measure based on pairwise orderings where pairs are counted if they are concordant and otherwise [18]. Since the proposed representation is based on comparisons, it does not require scaling or normalization and will work expression measurements obtained with different experimental technologies.

### 2.2 Survival Prediction

#### Problem Description and the Survival Model

We apply the PRER representation for the survival prediction problem. For each cancer type, the data is of the form, 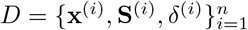; *n* is the number of patients. For each patient, **x** is the derived features from protein expression data, **S** is the overall survival time, and *δ* denotes censoring. We use random survival forests for the problem. Random Survival Forest(RSF)[19] is a non-parametric method and has been shown to perform well in survival prediction. It is an ensemble method wherein the base learner is a tree, and each tree is grown on a randomly drawn bootstrap sample. Furthermore, in growing a tree, a randomly selected subset of features is chosen as the candidate features for splitting at each node of the tree. The node is split with the feature among the candidate features that maximize survival difference between child nodes. We used the default values for the rfsrc package [19], where the number of trees is 1000, the number of random splits to consider for each candidate splitting variable is set to 10, and the default splitting rule for a node implements log-rank splitting [20, 21].

#### Molecular and Clinical Data

We test the method on ten different cancer types: ovarian adenocarcinoma (OV), breast invasive carcinoma (BRCA), glioblastoma multiforme (GBM), head and neck squamous cell carcinoma (HNSC), kidney renal clear cell carcinoma (KIRC), lung adenocarcinoma (LUAD), lung squamous cell carcinoma (LUSC), bladder urothelial carcinoma (BLCA), colon adenocarcinoma (COAD), uterine corpus endometrial carcinoma (UCEC). The number of patients for each cancer type range from 112 to 841, and the number of all patients is 3253. Details for each cancer are provided in Supplementary Table 1. For each cancer type, the number of patients is given at Supplementary Table 1. We obtained TCGA protein expression data and patient survival data from UCSC Cancer Browser (https://genome-cancer.ucsc.edu) (April 11, 2017). The protein expression is quantified by reverse-phase protein array (RPPA). There are measurements for 131 proteins, some include protein phosphorylated forms of the proteins.. For example, RPPA data include STAT3 and STAT3PY705, where STAT3 is Signal Transducer And Activator Of Transcription 3 protein, and STAT3PY705 is the phosphorylation of STAT3 at tyrosine 705 residue. We were able to map all proteins to the PPI network. Since PPIs do not represent phosphorylated forms separately, we use the unphosphorylated node when obtaining the neighborhood for the phosphorylated protein.

#### Protein-Protein Interaction Network

We obtained the protein-protein interaction (PPI) network from the InBioMap platform (April 11, 2017). InBioMap specifies a confidence score for each edge, representing the support of the interaction in the literature. The interactions that have lower than 0.1 confidence cut-off are eliminated from the network. The final network used in this study includes 17, 653 proteins and 625, 641 interactions between those proteins.

## 3 Results and Discussion

To understand if PRER representation captures the molecular expression profiles better than the individual protein expression values, we use these representations for survival prediction. We first build two sets of survival prediction models for the 10 cancer types. In building these two sets of models, only the feature representations differ. In the first one, we use the protein expression values as input, which is the typical approach taken in survival prediction. In contrast, in the second one, we use the proposed PRER representation.

Next, we compare our model with two competitive methods from the literature. The first model is by Hofree et al. [7], which uses network propagation to diffuse information on each protein. In their original method, Hofree et al. [7] uses mutation data for patient stratification. Here, we used the protein expression data, use the same network propagation method to diffuse the expression values over the network. We input the feature vector that contains the diffused feature values into RSF as patient features. We implemented this algorithm in R and set network propagation parameter *α* to 0.5 and run the RSF model with default parameters. As the second method, we use Reweighted RSF (RRSF) method, proposed by Wang and Liu [13]. RRSF weights the features in random sampling step of RSF model with their topological importance in the PPI network. For RRSF, we use the authors’ R implementation.

In all the models trained, we randomly split the samples into train and test groups: 80% as the training set and 20% as the test set. We train 100 such models in 100 test runs. In each of these models, we perform a univariate feature selection based on the hazard ratio of the Cox model [22] with the exception of the RRSF model. We use the p-values of the likelihood ratio test to quantify the significance of hazard ratio, and features with p-value ≤ 0.05 are retained for model training. With given random walk parameters in Section 2, and the InBioMap PPI, using 131 proteins in RPPA, PRER produces 1909 dimensional feature vectors for each patient. After applying univariate feature filtering with Cox model [22] to these 1909 dimensional feature vectors, and we provide the average number of features that pass the Cox screen step for each cancer type at Supplementary Table 2. Finally, the models are evaluated by the Concordance-Index (C-index) [23] on the test data. The pipeline of the model training and evaluation is summarized in Figure 2a.

**Fig. 2:**
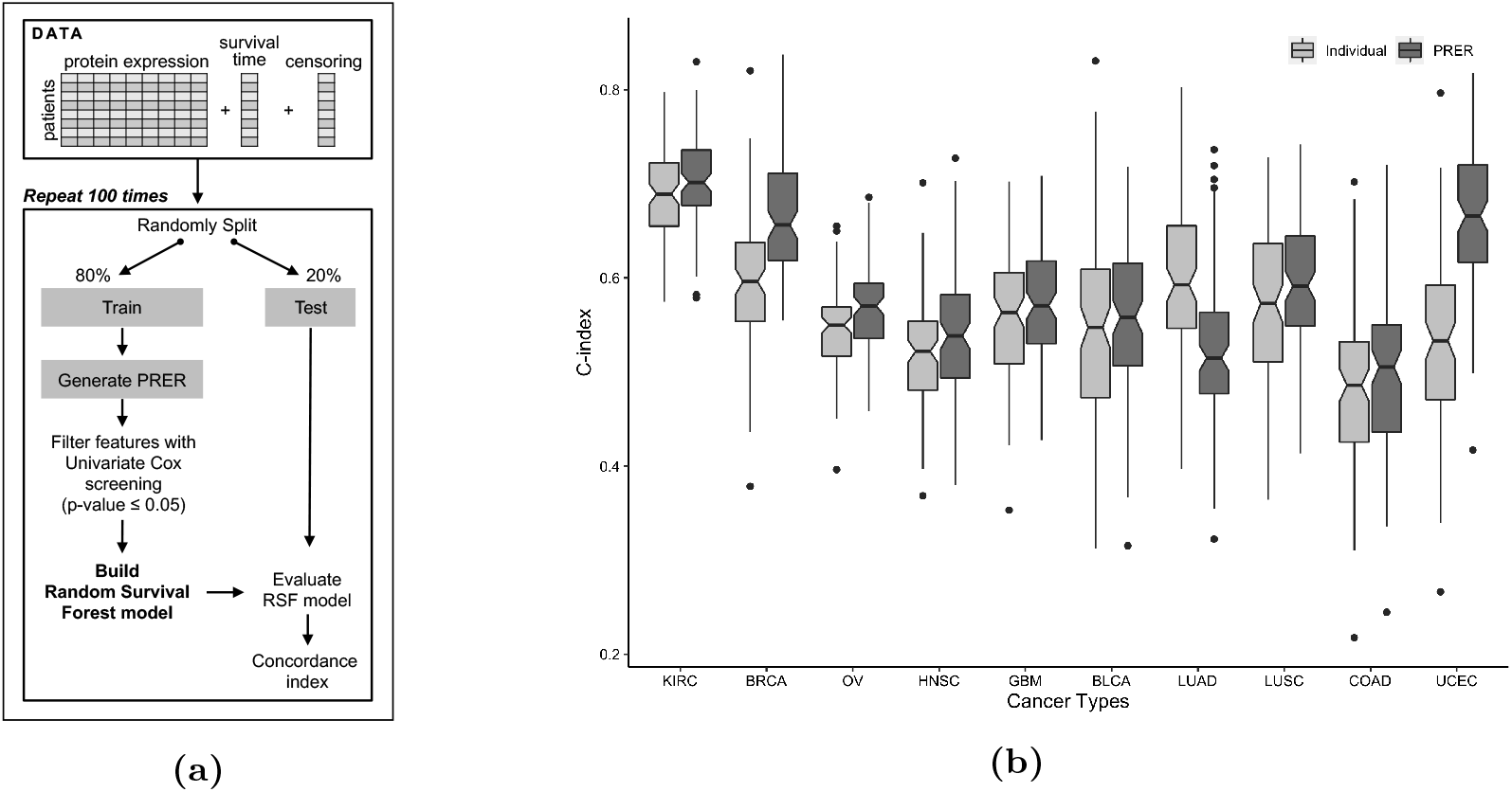
**(a)** The pipeline for survival prediction. The step that involves generating PRER is skipped when the experiment is run with the alternative method of individual expression values. **(b)** Comparison of RSF model performances that are trained with individual proteins and pairwise ranking representations for different cancer types. The distribution is over 100 models trained that have different random train and test splits. The performances of the models that use the individual expression values as features (Individual) and PRER representation as features (PRER) are compared in each case.

### 3.1 Survival Prediction Performance of PRER

We first compare PRER with the models trained with individual features. Figure 2b compares the distribution of C-indices for 100 models trained with the two different feature representations for 10 different cancer types. In 9 of 10 cancer types, PRER representation yields statistically significant improvements (Wilcoxon signed-rank test, (BH adjusted p-value < 0.05)). The C-index quantiles of 100 bootstrap results and corresponding *p-values* are listed in Supplementary Table 3. The best improvements are found in *UCEC*, *BRCA*, *KIRC* and *OV*.

Next, we compare PRER with two other competitive methods, network propogation by Hofree et al.[7] and RRSF by [13]. Supplementary Figure 1 and Tables 3 details these result of performance comparisons. To summarize the performance of PRER against the two competitor methods, we present a win/tie/loss table (Table 1). In this table, a win count corresponds to the number of cancer types on which PRER achieves statistically significant performance improvements. In contrast, the loss count denotes the number of cancer types on which the compared method achieves statistically significant improvements. If none of the methods can achieve a significant improvement compared to the other, we mark it as a tie. We observe that PRER outperforms the network propagation representation and the RSSF method in 5 of the cancers, ties with them in 4 cancer types and underperforms in one cancer. The cancer type that where PRER underperforms is LUAD, which we do not observe any improvement with PRER representation (Figure 2b).

**Table 1:** Win/Tie/Loss counts of **PRER** against competing methods. PRER is compared against each model over 100 trained models, where each model is trained on a different train/test split. The comparisons are based on one-sided Wilcoxon signed rank test with BH multiple hypothesis test correction at the significance level of 0.05. The Hofree et al. method is the network propogation algorithm [7]. RRSF stands for reweighted random survival algorithm by Wang and Liu [13].

|  | vs. Individual | vs. Hofree et al. | vs. RRSF |
| --- | --- | --- | --- |
| **PRER** | (9 / 0 / 1) | (5 / 4 / 1) | (5 / 4 / 1) |

### 3.2 Effect of Different Parameter Choices on PRER Performance

#### Effect of Choice of Protein-Protein Interaction Network

To understand the effect of PPI change, we repeat the experiments on 10 cancers using a second network. For this, we use the PPI network made available by the IntAct database [24]. This time, we found statistically significant improvements in 6 out of 10 cancer types. We provide C-index quantiles and Wilcoxon signed-rank test adjusted p-values in Supplementary Table 5. The difference between the two sets of results could be due to the edge density of differences of the networks. The InBioMap network contains 17, 653 nodes and 625, 641 edges whereas IntAct database contain 583, 756 edges and 29, 629 nodes. Although the number of nodes is higher in the IntAct PPI, the edge density of InBioMap is four times higher than that of IntAct’s (0.004 vs. 0.001). The edge density is calculated as the number of edges divided by the possible number of edges. This illustrates that PRER performance, as expected, is dependent on the PPI network used.

#### Effect of Random Walk Parameters

In PRER, we define the neighborhood of a protein using random walks. There are several input parameters for the random walk technique which we use: the number of walks, walk length, *p* and *q*. To see their influence on the output of PRER, we conduct runs with various choices of these parameters. Supplementary Figures 3 - 12 contains this parameter sensitivity results. For each cancer, the effect of a parameter is different. For example, as the number of walks or the length of the walk increases, the prediction performance slightly increases for BRCA and the GBM. However, we observe the opposite effect for BLCA and UCEC. For the other cancers, there is not much of a difference. The change in *p* and *q* does not drastically change the performance. These hyperparameters can be tuned for each cancer separately with larger patient cohorts. We provide an analysis of each parameter for different cancers in Supplementary Figures 3 - 12.

#### Effect of the Amount of Difference between the Protein Expression Levels

PRER representation assigns a binary value to a specific protein pair, either 1 or 1 based on pairwise comparison of the protein expression levels. We experimented with an alternative representation, for which we assign the feature value 0 if the difference between the expression values is less than 10% of the compared neighbor, and otherwise, we assign 1 and −1 based on the comparison. We call this representation as ternary PRER. We compare this ternary feature representation model against the binary feature representation model. We observe no improvements in eight cancer types, but we observe improvement in GBM (p-value = 0.01) and UCEC (p-value = 0.02). The detailed results are presented in Supplementary Table 4 and Supplementary Figure 2. What to consider as a meaningful expression difference between pairs of protein is probably depends on both the tissue and the protein pairs question. When compared matched normals are available, the difference threshold could be decided per protein pair and per cancer type. Since we did not have it here, we choose the method with the least assumption and rely on the RSF model to pick up the meaningful features. In future work with richer datasets, this step could be improved.

### 3.3 Predictive PRER Features

We seek to determine the features ranked as significant in the RSF models trained with PRER features. Note that in these models, pairs of proteins constitute the features. The importance of a particular feature is quantified by the performance difference between the models trained with the original feature vector and the case where the feature vector values are permuted [25]. A significant difference indicates a feature whose absence degrades the model performance. As there are 100 models trained on the repeatedly split data, we calculate the overall feature importance scores over these models as the sum of the scores. We show the normalized feature importance scores for ovarian cancer (OV) in Figure 3a. The feature importance scores for other cancer types are available in Supplementary Figures 13-21).

**Fig. 3:**
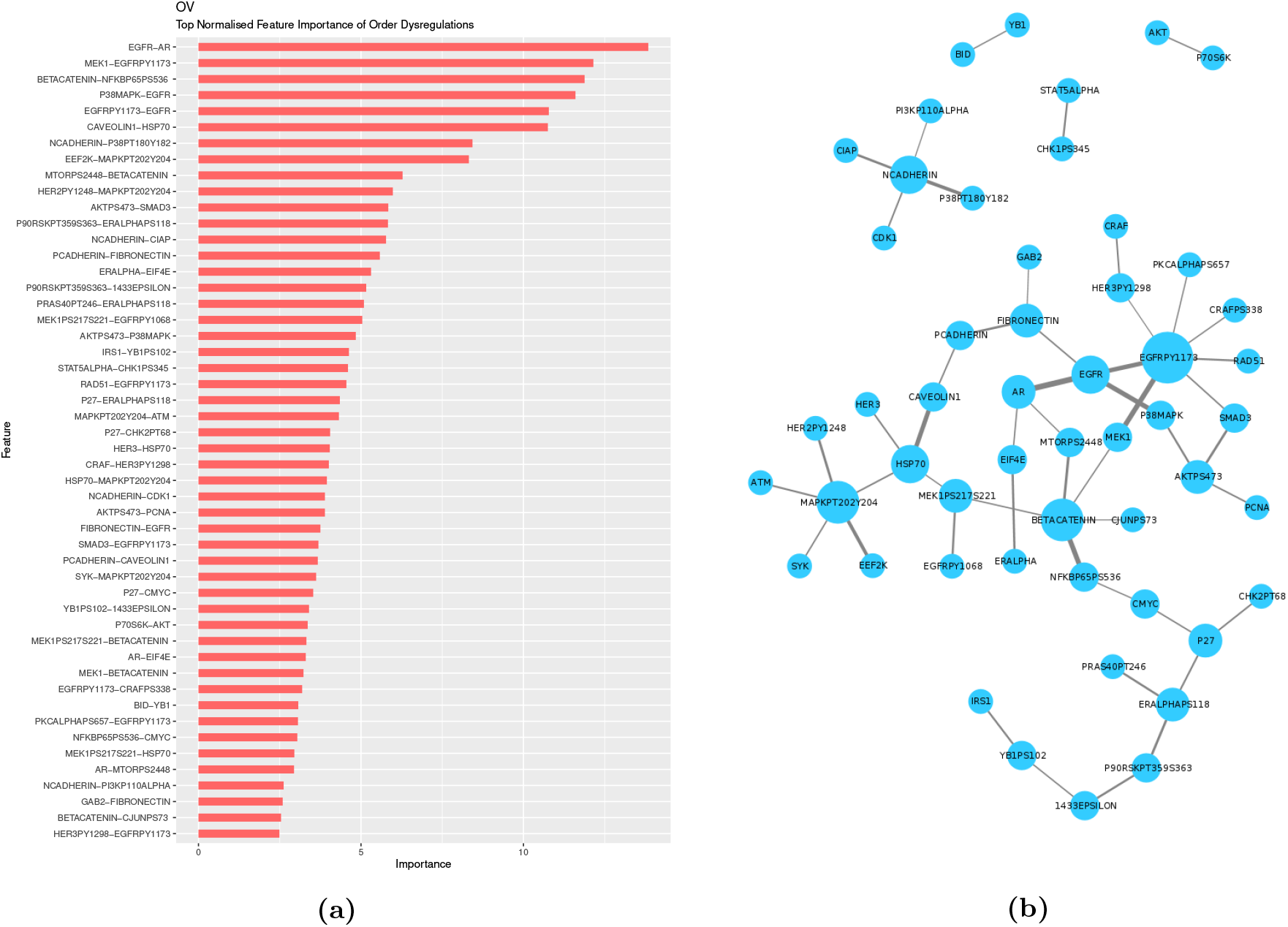
**(a)** The variable importance of significant pairwise ranking representations for ovarian cancer. **(b)** Nodes represent proteins that appear in the top 50 pairwise ranking representations for ovarian cancer; each edge indicate that two proteins participate in a pairwise rank order feature together. For cases where the expression value pertains to the phosphorylated state of the protein, the ids include the phosphosite’s residue position and the amino acid type of the phosphosite.

As shown in Figure 3a, some proteins repeatedly show up as partners in the list of important genes. To analyze these relationships, we form a network where the nodes represent proteins that participate in the top 50 PRER features. Edges are formed when a given protein pair is found to be partners in a PRER feature. Figure 3b demonstrates that some proteins emerge as important in many pairs. Several studies support these genes’ association with ovarian cancer. Epidermal growth factor receptor protein (EGFR) and its phosphorylated state EGFRPY1173 are among the top PRER features. EGFR is a receptor protein that receives and transmits signals from the environment to the cell and is the target of drugs in therapies for many cancer types, including ovarian cancer [26, 27]. Marozkina et al. provide results that changes in expression of EGFR may lead to ovarian carcinoma. Others [29, 30, 31] also claim that up-regulation of EGFR expression promotes ovarian cancer. Interestingly, Li et al. and Ilekis et al. demonstrate that the levels of EGFR and androgen receptor (AR), which constitute the top feature of PRER in Figure 3a, are interacted in ovarian cancer.

Another important protein that participates in important features is Caveolin-1 (CAV1). CAV1 takes on critical roles in cell survival, cell proliferation, cell migration and programmed cell death [33]. An earlier study by Wiechen et al. report that CAV1 is dysregulated among ovarian cancer patients based on microarray expression data. Others also report that CAV1 is reported to be dysregulated in different cancer types and its role in chemotherapy resistance [35, 36].

We list the top-ranked PRER pairs for each cancer in Table 2. We provide the Kaplan-Meier (KM) plots of the top feature for KIRC and BLCA based on overall survival in Figure 4. Based on only one feature, the patients can be grouped into groups that differ significantly in their survival distributions. We provide the KM plots of top-ranked features for the other cancers in Supplementary Figure 31. The confounding factors such as the age and sex of the patient may influence protein expressions. Therefore, we adjust survival curves for the confounding effects. We also apply log-rank tests to adjusted curves and see that age and sex adjustment gives the same p-values of top PRERs.

**Table 2:** The top PRER feature in each cancer type. The relative expression level of this feature is found to be important in the RSF model. The gene symbols of the corresponding gene are listed. The letter P after the gene symbol indicates that this is the phosphorylated version of the protein. The type of phosphosite and its residue number is provided.

| Cancer | Top Rank PRER Protein Pair |
| --- | --- |
| BLCA | NCADHERIN-SRCPY416 |
| BRCA | DVL3-P38MAPK |
| COAD | MRE11-HER3PY1298 |
| GBM | NF2-EGFR |
| HNSC | ECADHERIN-PAXILLIN |
| KIRC | 4EBP1T37T46-AR |
| LUAD | XRCC1-CYCLINB1 |
| LUSC | PAXILLIN-YAP |
| OV | EGFR-AR |
| UCEC | EIF4E-AKT |

**Fig. 4:**
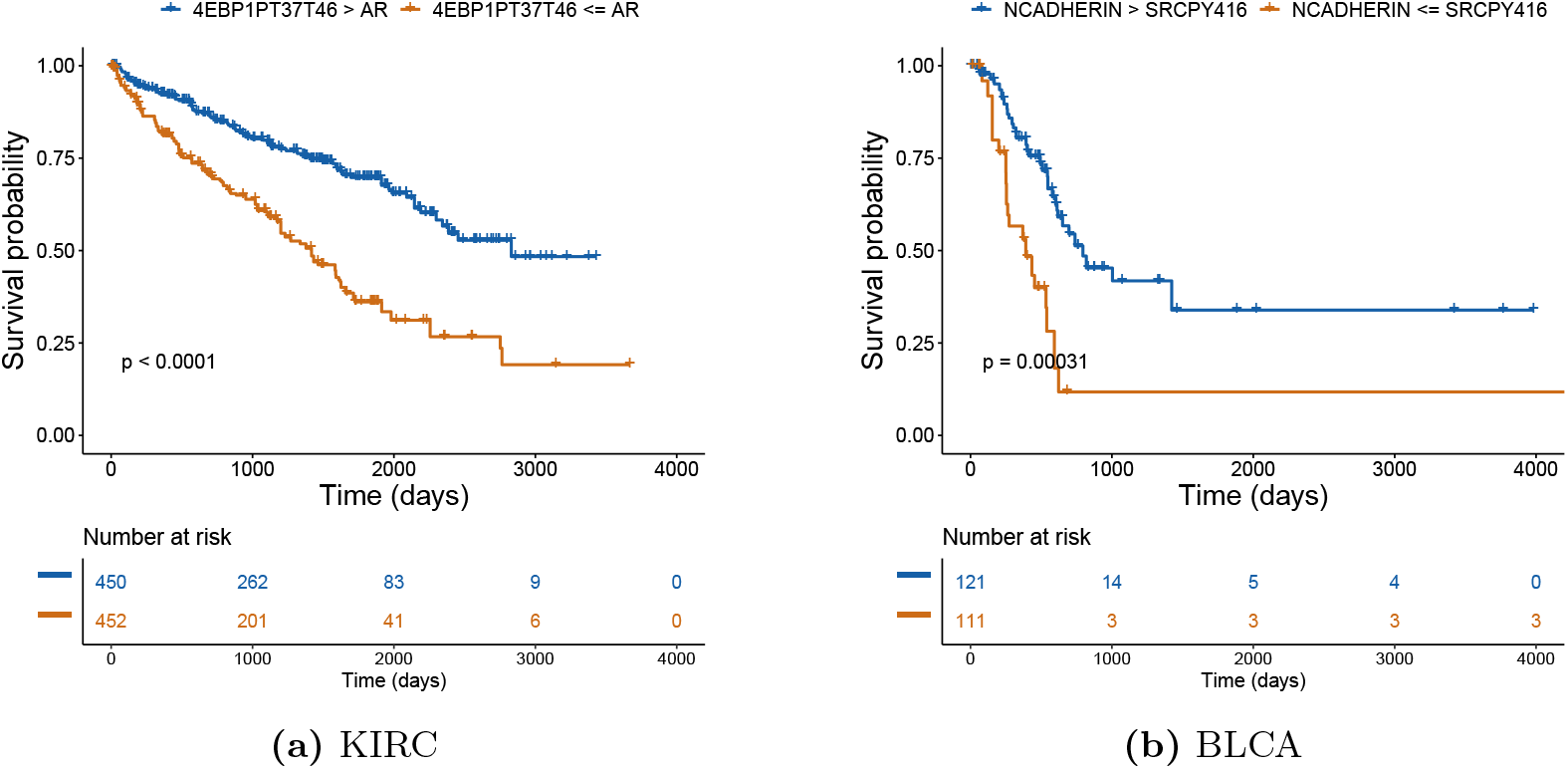
Age and sex adjusted Kaplan-Meier plots for a) KIRC and b) BLCA based on overall survival. Number at risk denotes the number of patients at risk at a given time, and p-value is calculated with the log-rank test.

We should note that many of the proteins reported in the RPPA assay in the TCGA study are selected due to their relevance to cancer. Thus, these important genes are likely to exhibit the individual importance of PRER partners. Therefore, we suggest an alternative way to exclusively analyze those features which emerge as important in the next section.

### 3.4 Proteins that Emerge as Important only in the PRER Representation

Since many of the proteins that are in the protein expression data are cancer-related, it is not surprising that they are found to be relevant to cancer. However, proteins that emerge as important in the PRER representation but are not highly ranked in the models trained with individual protein expression values would be interesting. These sets of proteins will reveal proteins whose relative expression states to their neighbors are important as opposed to the expression level being up or down-regulated. To identify these proteins, we first assign a feature importance score to each protein in the PRER representation. As the features are pairs of proteins in the PRER models, we calculate the feature importance of a protein by averaging the importance of the PRER feature importance in which this protein contributes. Let *f*_*i,j*_ denotes the feature importance score of the protein pair *i* and *j*. We calculate the individual feature importance score for molecule *i* as follows:

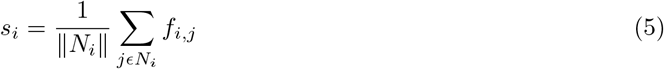

where *N*_*i*_ is the set of all pairwise ranking representations that include molecule *i*. *s*_*i*_ represents the average importance of molecule *i* concerning the expression levels of other proteins in its neighborhood. We get the rank order of each protein based on *s*_*i*_, and a lower rank indicates that the protein is important. Let *r*_*p*_ be the protein’s rank in the models with PRER representation and let *r*_*q*_ be the rank order in the models trained with individual protein expressions. To find the proteins whose ranks are low in the models trained with protein expression but are highly ranked in the PRER models, we measure the differences of feature ranks, *r*_*q*_ − *r*_*p*_. Table 3 lists the top 10 proteins in each cancer based on this *r*_*q*_ − *r*_*p*_ difference. We provide the full list of the ranks and differences in Supplementary Table 6. A large positive difference points to those proteins for which the relative expression relations of this protein to other proteins in its neighborhood carry prognostic value as opposed to its expression value.

**Table 3:**
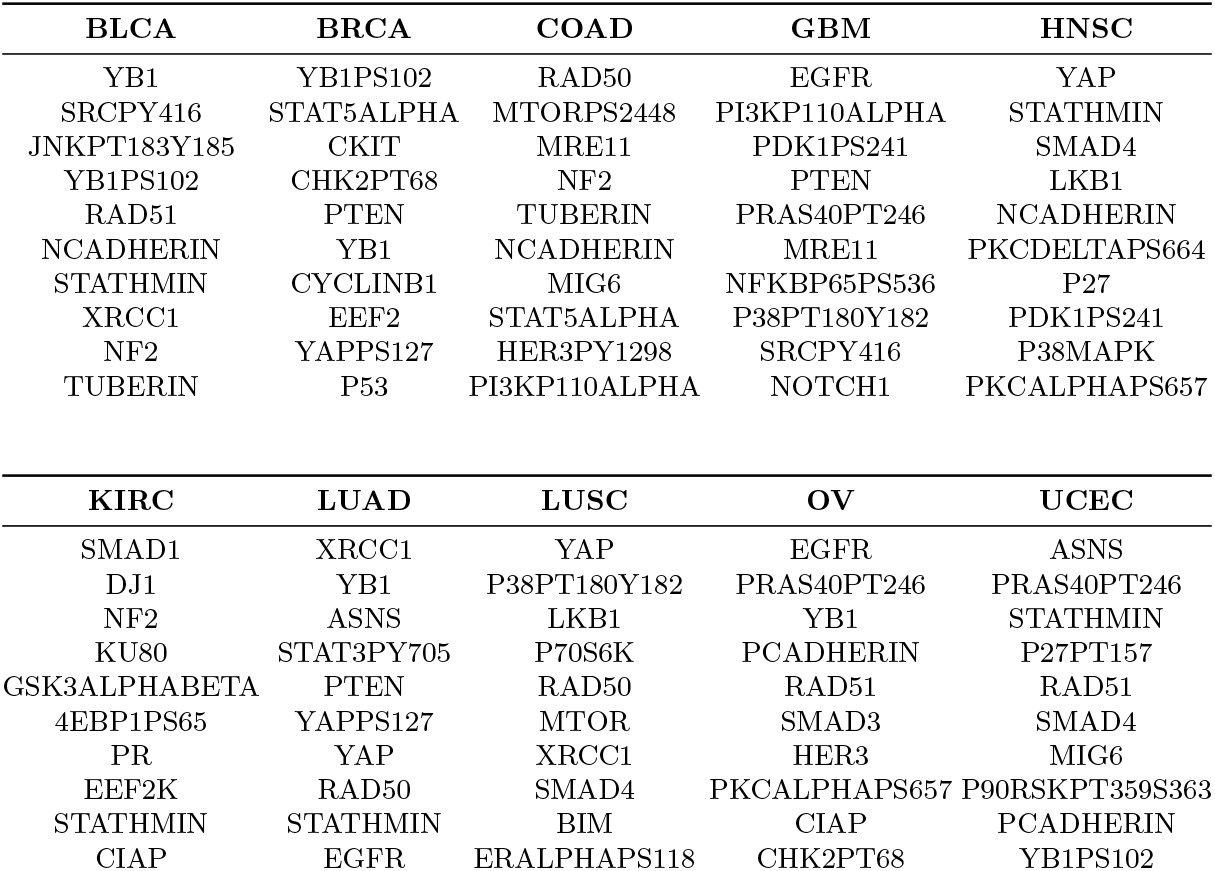
Top-10 rank differentiated features in each cancer with PRER.

We analyze a subset of the proteins in Table 3. The relevance of the relative expressions of proteins for survival is not reported. Some proteins that are known to be cancer drivers and perturbed in cancers such as PTEN or EGFR do not rank high in the model wherein the protein expression data is used as input, but in PRER models, they emerge as important. For example, EGFR is ranked as the 16^*th*^ most important feature for ovarian cancer in the models trained with PRER, while it is ranked as the least significant one in the models trained with individual expressions only. Similarly, for GBM, EGFR is ranked as the least significant protein in individual expression models, while it is ranked as the 13^*th*^ most significant feature in PRER. Thus, the PRER models actually highlight that the dysregulation of EGFR expression with respect to its neighbors is an important feature. Below we mention other interesting observations in Table 3.

STAT3PY705 (STAT3 phosphorylation at tyrosine 705), phosphorylated state of STAT3 (Signal Transducer and Activator of Transcription 3) protein, and STAT5ALPHA (Signal Transducer And Activator Of Transcription 5A) also appear in multiple cancer types. While we observe STAT3PY705 as significant in LUAD, STAT5ALPHA appears in BRCA and COAD in Table 3. Activation in the STAT family is reported, especially for STAT3 and STAT5, in several cancer cell lines including head and neck, breast, kidney, ovarian and colorectal[37, 38, 39, 40].

YAPPS127 and YAP proteins, which are encoded with the YAP1 (Yes-associated protein 1) gene, found important in BRCA, HNSC, LUAD, and LUSC cancer types in Table 3. YAP1 is involved in the Hippo signaling pathway that is associated with the growth, development and repair of the cells, and influences the survival of multiple cancers [41]. Poma et al. reports that 17 genes (out of 32) in the Hippo pathway have effects on survival in more than 20 different cancer types and conclude that YAP1 is relevant to the survival of head and neck carcinoma, hepatocellular, lung adenocarcinoma, gastric, pancreatic and colorectal cancers. Further, other studies also suggest that survival for different cancer types is associated with the expression level of YAP1 and its differential expression is considered as a biomarker for bladder urothelial carcinoma (BLCA) [43], breast invasive carcinoma (BRCA) [44, 45, 46, 47], ovarian serous cystadenocarcinoma (OV) [48, 49].

The upregulation of STATHMIN is linked with poor survival for primary HNSC [50], and Kouzu et al. suggest that it may be used for the prognosis and a therapeutic target for oral squamous-cell carcinoma, which is the most common type of HNSC. Likewise, the upregulation of STATHMIN is significantly correlated with several cancer types such as LUAD [52], gastric cancer [53, 54], UCEC [55], OV [56] and BRCA [57, 58, 59].

YB1 and its phosphorylated state YB1PS102 show correlation with many genes that have functions such as resistance to drugs, transcription and translation of cancerous cells [60]. Although the down-regulation of YB1 is found to be correlated with the reduction in progression, development of cell and programmed cell death at various cancer cells such as breast, colon, lung, prostate and pediatric glioblastoma by some studies [61, 62], there are studies [63, 64, 65, 66, 67] showing the association between overexpression of YB1 and different cancer types such as breast, colorectal, glioblastoma, lung, liver, ovarian cancers.

## 4 Conclusion and Future Work

Accurate prediction of clinical outcomes such as survival success remains to be a challenge for cancer patients. If achieved, it can guide the decision-making process for choosing optimal treatment and surveillance strategies among alternative options. Typically, clinical or pathological features such as the patient’s age, tumor stage or grade are employed to predict the clinical outcomes. With the advent of high-throughput technologies, molecular descriptions of the tumors for a large number of patients across many cancer types have become available. However, it remains a significant challenge to use this data due to the high level of genomic heterogeneity among patients. This study proposes a novel patient representation method, PRER. PRER is based on a pairwise comparison of a protein’s expression values with the other proteins in its neighborhood on the PPI network. In this way, the relative expression level patterns with respect to the proteins in their neighborhood can be captured.

We showcase PRER in the task of survival prediction for ten different cancer types. PRER with Random Survival Forest (RSF) model achieves statistically significant improvements compared to the models with individual expression values in 9 of the 10 cancer types. We also suggest ways to delineate the importance of proteins not through their individual up or down-regulation patterns but their relative expressions compared to their neighbors. Such an analysis can provide fundamental mechanistic insights into the studied diseases. The identified pairwise relations could also help design therapies to regulate the pairwise interaction as opposed to regulating the expression level of one protein.

One limitation of the current study is that we use a generic protein-protein interaction network, disregarding whether the protein is expressed in the given cancer type tissue. As tissue specific reliable PPI networks become available, we can improve the survival models by incorporating these.A second limitation is that in PRER we use a comparison based on the protein expression levels and assign the feature value 1 or −1 based on this difference. We also experiment with a ternary representation where we require this difference to be 10% of the expression level of the protein neighbor compared. These are, of course, ad-hoc choices. What constitutes a large enough difference depends on the tissue type and the protein pair in question. For certain pairs, large differences could be tolerated due to regulatory feedback mechanisms among genes or proteins performing similar functions, while for certain pairs of proteins minuscule differences can have a large impact on the cellular processes. The ideal scenario would be to decide this threshold based on expression values of the same protein pair in matched normal samples. Since we do not have it here we choose the method with the least number of assumptions and rely on the RSF model to pick up the predictive features. In future work, with the increasing availability of richer datasets, this step can be improved.

In this work, since we aim to assess the PRER representation power, we only use features related to expression. The survival model can be further improved with clinical features such as age, duration of the follow-up, and cancer stage. PRER representation can be used with other data types, such as mRNA expression and DNA methylation. However, we should note that the number of features increases quadratically with the size of the original features as each feature is compared with its neighboring proteins. In this case, a more stringent feature filtering step, reducing the number of neighbors or a regularized prediction model will be helpful.

## 5 Acknowledgements

Authors thank Bilkent and Sabanci Universities internal funds. OT acknowledges support from Science Academy of Turkey BAGEP program. The results here are in part based upon data generated by the TCGA Research Network.

## Bibliography

[1] Y. Yuan, E. M. Van Allen, L. Omberg, N. Wagle, A. Amin-Mansour, A. Sokolov, L. A. Byers, Y. Xu, K. R. Hess, L. Diao et al., “Assessing the clinical utility of cancer genomic and proteomic data across tumor types,” Nature biotechnology, vol. 32, no. 7, p. 644, 2014.

[2] Z. Jagga and D. Gupta, “Classification models for clear cell renal carcinoma stage progression, based on tumor rnaseq expression trained supervised machine learning algorithms,” in BMC proceedings, vol. 8, no. S6. Springer, 2014, p. S2

[3] Z. Ding, S. Zu, and J. Gu, “Evaluating the molecule-based prediction of clinical drug responses in cancer,” Bioinformatics, vol. 32, no. 19, pp. 2891–2895, 2016.

[4] C. Suphavilai, D. Bertrand, and N. Nagarajan, “Predicting cancer drug response using a recommender system,” Bioinformatics, vol. 34, no. 22, pp. 3907–3914, 2018.

[5] L. Cowen, T. Ideker, B. J. Raphael, and R. Sharan, “Network propagation: a universal amplifier of genetic associations,” Nature Reviews Genetics, vol. 18, no. 9, p. 551, 2017.

[6] A.-L. Barabási, N. Gulbahce, and J. Loscalzo, “Network medicine: a network-based approach to human disease,” Nature reviews genetics, vol. 12, no. 1, pp. 56–68, 2011.

[7] M. Hofree, J. P. Shen, H. Carter, A. Gross, and T. Ideker, “Network-based stratification of tumor mutations,” Nature methods, vol. 10, no. 11, pp. 1108–1115, 2013.

[8] H.-Y. Chuang, E. Lee, Y.-T. Liu, D. Lee, and T. Ideker, “Network-based classification of breast cancer metastasis,” Molecular systems biology, vol. 3, no. 1, 2007.

[9] I. W. Taylor, R. Linding, D. Warde-Farley, Y. Liu, C. Pesquita, D. Faria, S. Bull, T. Pawson, Q. Morris, and J. L. Wrana, “Dynamic modularity in protein interaction networks predicts breast cancer outcome,” Nature biotechnology, vol. 27, no. 2, p. 199, 2009.

[10] A. P. Crijns, R. S. Fehrmann, S. de Jong, F. Gerbens, G. J. Meersma, H. G. Klip, H. Hollema, R. M. Hofstra, G. J. te Meerman, E. G. de Vries et al., “Survival-related profile, pathways, and transcription factors in ovarian cancer,” PLoS medicine, vol. 6, no. 2, p. e1000024, 2009.

[11] W. Zhang, T. Ota, V. Shridhar, J. Chien, B. Wu, and R. Kuang, “Network-based survival analysis reveals subnetwork signatures for predicting outcomes of ovarian cancer treatment,” PLoS Comput Biol, vol. 9, no. 3, p. e1002975, 2013.

[12] C. Winter, G. Kristiansen, S. Kersting, J. Roy, D. Aust, T. Knösel, P. Rümmele, B. Jahnke, V. Hentrich, F. Rückert et al., “Google goes cancer: improving outcome prediction for cancer patients by network-based ranking of marker genes,” PLoS computational biology, vol. 8, no. 5, p. e1002511, 2012.

[13] W. Wang and W. Liu, “Integration of gene interaction information into a reweighted random survival forest approach for accurate survival prediction and survival biomarker discovery,” Scientific reports, vol. 8, no. 1, pp. 1–14, 2018.

[14] D. Geman, C. d’Avignon, D. Q. Naiman, and R. L. Winslow, “Classifying gene expression profiles from pairwise mrna comparisons,” Statistical applications in genetics and molecular biology, vol. 3, no. 1, pp. 1–19, 2004.

[15] A. Magen, A. D. Sahu, J. S. Lee, M. Sharmin, A. Lugo, J. S. Gutkind, A. A. Scháffer, E. Ruppin, and S. Hannenhalli, “Beyond synthetic lethality: Charting the landscape of pairwise gene expression states associated with survival in cancer,” Cell reports, vol. 28, no. 4, pp. 938–948, 2019.

[16] J. N. Weinstein, E. A. Collisson, G. B. Mills, K. R. M. Shaw, B. A. Ozenberger, K. Ellrott, I. Shmulevich, C. Sander, J. M. Stuart, C. G. A. R. Network et al., “The cancer genome atlas pan-cancer analysis project,” Nature genetics, vol. 45, no. 10, p. 1113, 2013.

[17] A. Grover and J. Leskovec, “node2vec: Scalable feature learning for networks,” KDD : proceedings. International Conference on Knowledge Discovery & Data Mining, vol. 2016, pp. 855–864, 2016.

[18] M. G. Kendall, “A new measure of rank correlation,” Biometrika, vol. 30, no. 1/2, pp. 81–93, 1938.

[19] H. Ishwaran, U. B. Kogalur, E. H. Blackstone, and M. S. Lauer, “Random survival forests,” The Annals of Applied Statistics, pp. 841–860, 2008.

[20] M. R. Segal, “Regression trees for censored data,” Biometrics, pp. 35–47, 1988.

[21] M. LeBlanc and J. Crowley, “Survival trees by goodness of split,” Journal of the American Statistical Association, vol. 88, no. 422, pp. 457–467, 1993.

[22] T. Therneau, “A package for survival analysis in s. r package version 2.37-4. 2013,” 2013.

[23] F. E. Harrell Jr, R. M. Califf, D. B. Pryor, K. L. Lee, R. A. Rosati et al., “Evaluating the yield of medical tests,” Jama, vol. 247, no. 18, pp. 2543–2546, 1982.

[24] S. Kerrien, B. Aranda, L. Breuza, A. Bridge, F. Broackes-Carter, C. Chen, M. Duesbury, M. Dumousseau, M. Feuermann, U. Hinz et al., “The intact molecular interaction database in 2012,” Nucleic acids research, vol. 40, no. D1, pp. D841–D846, 2012.

[25] H. Ishwaran et al., “Variable importance in binary regression trees and forests,” Electronic Journal of Statistics, vol. 1, pp. 519–537, 2007.

[26] L. G. Hudson, R. Zeineldin, M. Silberberg, and M. S. Stack, “Activated epidermal growth factor receptor in ovarian cancer,” in Ovarian Cancer. Springer, 2009, pp. 203–226.

[27] J. A. Wilken, T. Badri, S. Cross, R. Raji, A. D. Santin, P. Schwartz, A. J. Branscum, A. T. Baron, A. I. Sakhitab, and N. J. Maihle, “Egfr/her-targeted therapeutics in ovarian cancer,” Future medicinal chemistry, vol. 4, no. 4, pp. 447–469, 2012.

[28] N. V. Marozkina, S. M. Stiefel, H. F. Frierson Jr, and S. J. Parsons, “Mmtv-egf receptor transgene promotes preneoplastic conversion of multiple steroid hormone-responsive tissues,” Journal of cellular biochemistry, vol. 103, no. 6, pp. 2010–2018, 2008.

[29] I. Dimova, B. Zaharieva, S. Raitcheva, R. Dimitrov, N. Doganov, and D. Toncheva, “Tissue microarray analysis of egfr and erbb2 copy number changes in ovarian tumors,” International Journal of Gynecological Cancer, vol. 16, no. 1, pp. 145–151, 2006.

[30] J. V. Ilekis, J. P. Connor, G. S. Prins, K. Ferrer, C. Niederberger, and B. Scoccia, “Expression of epidermal growth factor and androgen receptors in ovarian cancer,” Gynecologic oncology, vol. 66, no. 2, pp. 250–254, 1997.

[31] I. Skirnisdóttir, B. Sorbe, and T. Seidal, “The growth factor receptors her-2/neu and egfr, their relationship, and their effects on the prognosis in early stage (figo i-ii) epithelial ovarian carcinoma,” International Journal of Gynecological Cancer, vol. 11, no. 2, pp. 119–129, 2001.

[32] A. J. Li, D. R. Scoles, K. U. Armstrong, and B. Y. Karlan, “Androgen receptor cytosine-adenine-guanine repeat polymorphisms modulate egfr signaling in epithelial ovarian carcinomas,” Gynecologic oncology, vol. 109, no. 2, pp. 220–225, 2008.

[33] C. Boscher and I. R. Nabi, “Caveolin-1: role in cell signaling,” in Caveolins and Caveolae. Springer, 2012, pp. 29–50.

[34] K. Wiechen, L. Diatchenko, A. Agoulnik, K. M. Scharff, H. Schober, K. Arlt, B. Zhumabayeva, P. D. Siebert, M. Dietel, R. Scháfer et al., “Caveolin-1 is down-regulated in human ovarian carcinoma and acts as a candidate tumor suppressor gene,” The American journal of pathology, vol. 159, no. 5, pp. 1635–1643, 2001.

[35] L. A. Carver and J. E. Schnitzer, “Caveolae: mining little caves for new cancer targets,” Nature Reviews Cancer, vol. 3, no. 8, pp. 571–581, 2003.

[36] M. Zhang and S. Luo, “Gene expression profiling of epithelial ovarian cancer reveals key genes and pathways associated with chemotherapy resistance,” Genet Mol Res, vol. 15, no. 1, p. 11, 2016.

[37] R. Buettner, L. B. Mora, and R. Jove, “Activated stat signaling in human tumors provides novel molecular targets for therapeutic intervention,” Clinical cancer research, vol. 8, no. 4, pp. 945–954, 2002.

[38] H. Yu and R. Jove, “The stats of cancernew molecular targets come of age,” Nature Reviews Cancer, vol. 4, no. 2, p. 97, 2004.

[39] A. Lavecchia, C. Di Giovanni, and E. Novellino, “Stat-3 inhibitors: state of the art and new horizons for cancer treatment,” Current medicinal chemistry, vol. 18, no. 16, pp. 2359–2375, 2011.

[40] I. Souissi, I. Najjar, L. Ah-Koon, P. O. Schischmanoff, D. Lesage, S. Le Coquil, C. Roger, I. Dusanter-Fourt, N. Varin-Blank, A. Cao et al., “A stat3-decoy oligonucleotide induces cell death in a human colorectal carcinoma cell line by blocking nuclear transfer of stat3 and stat3-bound nf-*κ*b,” BMC cell biology, vol. 12, no. 1, p. 14, 2011.

[41] E. Lorenzetto, M. Brenca, M. Boeri, C. Verri, E. Piccinin, P. Gasparini, F. Facchinetti, S. Rossi, G. Salvatore, M. Massimino et al., “Yap1 acts as oncogenic target of 11q22 amplification in multiple cancer subtypes,” Oncotarget, vol. 5, no. 9, p. 2608, 2014.

[42] A. M. Poma, L. Torregrossa, R. Bruno, F. Basolo, and G. Fontanini, “Hippo pathway affects survival of cancer patients: extensive analysis of tcga data and review of literature,” Scientific reports, vol. 8, no. 1, p. 10623, 2018.

[43] J.-Y. Liu, Y.-H. Li, H.-X. Lin, Y.-J. Liao, S.-J. Mai, Z.-W. Liu, Z.-L. Zhang, L.-J. Jiang, J.-X. Zhang, H.-F. Kung et al., “Overexpression of yap 1 contributes to progressive features and poor prognosis of human urothelial carcinoma of the bladder,” BMC cancer, vol. 13, no. 1, p. 349, 2013.

[44] F. Cheng, J. Zhao, A. B. Hanker, M. R. Brewer, C. L. Arteaga, and Z. Zhao, “Transcriptome-and proteome-oriented identification of dysregulated eif4g, stat3, and hippo pathways altered by pik3ca h1047r in her2/er-positive breast cancer,” Breast cancer research and treatment, vol. 160, no. 3, pp. 457–474, 2016.

[45] L. Cao, P.-L. Sun, M. Yao, M. Jia, and H. Gao, “Expression of yes-associated protein (yap) and its clinical significance in breast cancer tissues,” Human pathology, vol. 68, pp. 166–174, 2017.

[46] S. K. Kim, W. H. Jung, and J. S. Koo, “Yes-associated protein (yap) is differentially expressed in tumor and stroma according to the molecular subtype of breast cancer,” International journal of clinical and experimental pathology, vol. 7, no. 6, p. 3224, 2014.

[47] H. M. Kim, W. H. Jung, and J. S. Koo, “Expression of yes-associated protein (yap) in metastatic breast cancer,” International journal of clinical and experimental pathology, vol. 8, no. 9, p. 11248, 2015.

[48] C. He, X. Lv, G. Hua, S. M. Lele, S. Remmenga, J. Dong, J. S. Davis, and C. Wang, “Yap forms autocrine loops with the erbb pathway to regulate ovarian cancer initiation and progression,” Oncogene, vol. 34, no. 50, p. 6040, 2015.

[49] Y. Xia, T. Chang, Y. Wang, Y. Liu, W. Li, M. Li, and H.-Y. Fan, “Yap promotes ovarian cancer cell tumorigenesis and is indicative of a poor prognosis for ovarian cancer patients,” PloS one, vol. 9, no. 3, p. e91770, 2014.

[50] H. Wu, W.-W. Deng, L.-L. Yang, W.-F. Zhang, and Z.-J. Sun, “Expression and phosphorylation of stathmin 1 indicate poor survival in head and neck squamous cell carcinoma and associate with immune suppression,” Biomarkers in medicine, vol. 12, no. 7, pp. 759–769, 2018.

[51] Y. Kouzu, K. Uzawa, H. Koike, K. Saito, D. Nakashima, M. Higo, Y. Endo, A. Kasamatsu, M. Shiiba, H. Bukawa et al., “Overexpression of stathmin in oral squamous-cell carcinoma: correlation with tumour progression and poor prognosis,” British journal of cancer, vol. 94, no. 5, pp. 717–723, 2006.

[52] L. Yurong, R. Biaoxue, L. Wei, M. Zongjuan, S. Hongyang, F. Ping, G. Wenlong, Y. Shuanying, and L. Zongfang, “Stathmin overexpression is associated with growth, invasion and metastasis of lung adenocarcinoma,” Oncotarget, vol. 8, no. 16, p. 26000, 2017.

[53] T. Jeon, M. Han, Y. Lee, Y. Lee, G. Kim, G. Song, G. Hur, J. Kim, H. Kim, S. Yoon et al., “Overexpression of stathmin1 in the diffuse type of gastric cancer and its roles in proliferation and migration of gastric cancer cells,” British journal of cancer, vol. 102, no. 4, pp. 710–718, 2010.

[54] X. Liu, H. Liu, J. Liang, B. Yin, J. Xiao, J. Li, D. Feng, and Y. Li, “Stathmin is a potential molecular marker and target for the treatment of gastric cancer,” International journal of clinical and experimental medicine, vol. 8, no. 4, p. 6502, 2015.

[55] W. Xi, W. Rui, L. Fang, D. Ke, G. Ping, and Z. Hui-Zhong, “Expression of stathmin/op18 as a significant prognostic factor for cervical carcinoma patients,” Journal of cancer research and clinical oncology, vol. 135, no. 6, pp. 837–846, 2009.

[56] D. Su, S. M. Smith, M. Preti, P. Schwartz, T. J. Rutherford, G. Menato, S. Danese, S. Ma, H. Yu, and D. Katsaros, “Stathmin and tubulin expression and survival of ovarian cancer patients receiving platinum treatment with and without paclitaxel,” Cancer, vol. 115, no. 11, pp. 2453–2463, 2009.

[57] L. H. Saal, P. Johansson, K. Holm, S. K. Gruvberger-Saal, Q.-B. She, M. Maurer, S. Koujak, A. A. Ferrando, P. Malmstróm, L. Memeo et al., “Poor prognosis in carcinoma is associated with a gene expression signature of aberrant pten tumor suppressor pathway activity,” Proceedings of the National Academy of Sciences, vol. 104, no. 18, pp. 7564–7569, 2007.

[58] G. Brattsand, “Correlation of oncoprotein 18/stathmin expression in human breast cancer with established prognostic factors,” British journal of cancer, vol. 83, no. 3, pp. 311–318, 2000.

[59] R. Golouh, T. Cufer, A. Sadikov, P. Nussdorfer, P. A. Usher, N. Brünner, M. Schmitt, R. Lesche, S. Maier, M. Timmermans et al., “The prognostic value of stathmin-1, s100a2, and syk proteins in er-positive primary breast cancer patients treated with adjuvant tamoxifen monotherapy: an immunohistochemical study,” Breast cancer research and treatment, vol. 110, no. 2, pp. 317–326, 2008.

[60] Y. Basaki, K.-i. Taguchi, H. Izumi, Y. Murakami, T. Kubo, F. Hosoi, K. Watari, K. Nakano, H. Kawaguchi, S. Ohno et al., “Y-box binding protein-1 (yb-1) promotes cell cycle progression through cdc6-dependent pathway in human cancer cells,” European journal of cancer, vol. 46, no. 5, pp. 954–965, 2010.

[61] Y. Basaki, F. Hosoi, Y. Oda, A. Fotovati, Y. Maruyama, S. Oie, M. Ono, H. Izumi, K. Kohno, K. Sakai et al., “Akt-dependent nuclear localization of y-box-binding protein 1 in acquisition of malignant characteristics by human ovarian cancer cells,” Oncogene, vol. 26, no. 19, p. 2736, 2007.

[62] A. Lasham, W. Samuel, H. Cao, R. Patel, R. Mehta, J. L. Stern, G. Reid, A. G. Woolley, L. D. Miller, M. A. Black et al., “Yb-1, the e2f pathway, and regulation of tumor cell growth,” Journal of the National Cancer Institute, vol. 104, no. 2, pp. 133–146, 2011.

[63] R. C. Bargou, K. Jürchott, C. Wagener, S. Bergmann, S. Metzner, K. Bommert, M. Y. Mapara, K.-J. Winzer, M. Dietel, B. Dörken et al., “Nuclear localization and increased levels of transcription factor yb-1 in primary human breast cancers are associated with intrinsic mdr1 gene expression,” Nature medicine, vol. 3, no. 4, p. 447, 1997.

[64] T. Kamura, H. Yahata, S. Amada, S. Ogawa, T. Sonoda, H. Kobayashi, M. Mitsumoto, K. Kohno, M. Kuwano, and H. Nakano, “Is nuclear expression of y box-binding protein-1 a new prognostic factor in ovarian serous adenocarcinoma?” Cancer: Interdisciplinary International Journal of the American Cancer Society, vol. 85, no. 11, pp. 2450–2454, 1999.

[65] K. Shibao, H. Takano, Y. Nakayama, K. Okazaki, N. Nagata, H. Izumi, T. Uchiumi, M. Kuwano, K. Kohno, and H. Itoh, “Enhanced coexpression of yb-1 and dna topoisomerase ii *α* genes in human colorectal carcinomas,” International journal of cancer, vol. 83, no. 6, pp. 732–737, 1999.

[66] K. Shibahara, K. Sugio, T. Osaki, T. Uchiumi, Y. Maehara, K. Kohno, K. Yasumoto, K. Sugimachi, and M. Kuwano, “Nuclear expression of the y-box binding protein, yb-1, as a novel marker of disease progression in non-small cell lung cancer,” Clinical cancer research, vol. 7, no. 10, pp. 3151–3155, 2001.

[67] M. Yasen, K. Kajino, S. Kano, H. Tobita, J. Yamamoto, T. Uchiumi, S. Kon, M. Maeda, G. Obulhasim, S. Arii et al., “The up-regulation of y-box binding proteins (dna binding protein a and y-box binding protein-1) as prognostic markers of hepatocellular carcinoma,” Clinical cancer research, vol. 11, no. 20, pp. 7354–7361, 2005.

