## Supplementary Document for "PRER: A Patient Representation with Pairwise Relative Expression of Proteins on Biological Networks"

### Supplementary Material for PRER: A Patient Representation with Pairwise Relative Expressions on Biological Networks

#### 1 Supplementary Tables

##### 1.1 Clinical and Molecular Patient Data

| **Number of Patients** | | |
|---|---|---|
| | censored | deceased |
| KIRC | 297 | 156 |
| OV | 194 | 213 |
| BRCA | 628 | 99 |
| LUAD | 147 | 72 |
| LUSC | 106 | 78 |
| COAD | 260 | 60 |
| HNSC | 98 | 114 |
| BLCA | 70 | 52 |
| UCEC | 347 | 50 |
| GBM | 73 | 139 |

**Table 1:** Number of censored and deceased patients for each cancer type.

| **Cancer Type** | **Average number of PRER features remain after Cox-fitering** |
|---|---|
| **BRCA** | 118 |
| **BLCA** | 336 |
| **COAD** | 114 |
| **HNSC** | 163 |
| **GBM** | 133 |
| **KIRC** | 778 |
| **LUAD** | 90 |
| **LUSC** | 189 |
| **OV** | 202 |
| **UCEC** | 312 |

**Table 2:** The average number of PRER features that pass the Cox filtering for each cancer.

#### 1.2 Prediction Performance

**Random Survival Forest C-indices**

| | C-index | | | | | p-value | BH adjusted p-value |
|---|---|---|---|---|---|---|---|
| | $1^{st}$ | $2^{nd}$ | median | $3^{rd}$ | $4^{th}$ | | |
| KIRC | 0.57 | 0.66 | 0.69 | 0.72 | 0.80 | | |
| KIRC-prer | 0.58 | 0.68 | **0.70** | 0.74 | 0.83 | $\mathbf{7.1x10^{-11}}$ | $\mathbf{2.4x10^{-10}}$ |
| KIRC-rrsf | 0.48 | 0.61 | 0.64 | 0.68 | 0.74 | 1 | 1 |
| KIRC-hofree | 0.56 | 0.67 | **0.70** | 0.73 | 0.80 | $\mathbf{7.8x10^{-4}}$ | $\mathbf{1.9x10^{-3}}$ |
| OV | 0.40 | 0.52 | 0.55 | 0.57 | 0.66 | | |
| OV-prer | 0.46 | 0.54 | **0.57** | 0.59 | 0.69 | $\mathbf{5.4x10^{-7}}$ | $\mathbf{1.3x10^{-6}}$ |
| OV-rrsf | 0.42 | 0.50 | 0.52 | 0.56 | 0.66 | 0.99 | 1 |
| OV-hofree | 0.41 | 0.53 | **0.56** | 0.59 | 0.68 | $\mathbf{2.7x10^{-4}}$ | $\mathbf{9.1x10^{-4}}$ |
| BRCA | 0.38 | 0.55 | 0.60 | 0.64 | 0.82 | | |
| BRCA-prer | 0.55 | 0.62 | **0.66** | 0.71 | 0.84 | $\mathbf{9.3x10^{-15}}$ | $\mathbf{4.7x10^{-14}}$ |
| BRCA-rrsf | 0.41 | 0.57 | **0.61** | 0.66 | 0.76 | 0.18 | 0.36 |
| BRCA-hofree | 0.39 | 0.54 | 0.59 | 0.63 | 0.77 | 1 | 1 |
| LUSC | 0.36 | 0.51 | 0.57 | 0.64 | 0.73 | | |
| LUSC-prer | 0.41 | 0.55 | **0.59** | 0.64 | 0.74 | $\mathbf{4.1x10^{-3}}$ | $\mathbf{6.8x10^{-3}}$ |
| LUSC-rrsf | 0.42 | 0.54 | **0.60** | 0.64 | 0.78 | $5.2x10^{-2}$ | 0.13 |
| LUSC-hofree | 0.34 | 0.55 | **0.60** | 0.66 | 0.75 | $\mathbf{1.8x10^{-4}}$ | $\mathbf{9x10^{-4}}$ |
| COAD | 0.22 | 0.43 | 0.49 | 0.53 | 0.70 | | |
| COAD-prer | 0.24 | 0.44 | **0.51** | 0.55 | 0.72 | **0.02** | **0.02** |
| COAD-rrsf | 0.28 | 0.47 | **0.53** | 0.61 | 0.75 | $\mathbf{1.1x10^{-4}}$ | $\mathbf{3.8x10^{-4}}$ |
| COAD-hofree | 0.29 | 0.45 | **0.53** | 0.58 | 0.72 | $\mathbf{2.8x10^{-3}}$ | $\mathbf{4.7x10^{-3}}$ |
| HNSC | 0.37 | 0.48 | 0.52 | 0.55 | 0.70 | | |
| HNSC-prer | 0.38 | 0.49 | **0.54** | 0.58 | 0.73 | $\mathbf{1.3x10^{-4}}$ | $\mathbf{2.7x10^{-4}}$ |
| HNSC-rrsf | 0.38 | 0.51 | **0.55** | 0.60 | 0.68 | $\mathbf{6.8x10^{-5}}$ | $\mathbf{3.4x10^{-4}}$ |
| HNSC-hofree | 0.36 | 0.50 | **0.54** | 0.57 | 0.76 | $\mathbf{1.2x10^{-3}}$ | $\mathbf{2.5x10^{-3}}$ |
| UCEC | 0.27 | 0.47 | 0.53 | 0.59 | 0.80 | | |
| UCEC-prer | 0.42 | 0.62 | **0.67** | 0.72 | 0.85 | $\mathbf{8.1x10^{-18}}$ | $\mathbf{8.1x10^{-17}}$ |
| UCEC-rrsf | 0.35 | 0.56 | **0.61** | 0.66 | 0.88 | $\mathbf{1.3x10^{-6}}$ | $\mathbf{1.3x10^{-5}}$ |
| UCEC-hofree | 0.30 | 0.52 | **0.56** | 0.63 | 0.73 | $\mathbf{5.8x10^{-5}}$ | $\mathbf{5.8x10^{-4}}$ |
| GBM | 0.35 | 0.51 | 0.56 | 0.61 | 0.70 | | |
| GBM-prer | 0.43 | 0.53 | **0.57** | 0.62 | 0.71 | **0.02** | **0.03** |
| GBM-rrsf | 0.41 | 0.52 | 0.55 | 0.60 | 0.69 | 0.58 | 0.97 |
| GBM-hofree | 0.36 | 0.52 | **0.56** | 0.61 | 0.75 | 0.13 | 0.19 |
| BLCA | 0.31 | 0.47 | 0.55 | 0.61 | 0.83 | | |
| BLCA-prer | 0.32 | 0.51 | **0.56** | 0.62 | 0.72 | **0.02** | **0.03** |
| BLCA-rrsf | 0.29 | 0.46 | 0.52 | 0.58 | 0.69 | 0.90 | 1 |
| BLCA-hofree | 0.24 | 0.50 | 0.54 | 0.59 | 0.70 | 0.86 | 1 |
| LUAD | 0.40 | 0.55 | 0.59 | 0.66 | 0.80 | | |
| LUAD-prer | 0.32 | 0.48 | 0.51 | 0.56 | 0.74 | 1 | 1 |
| LUAD-rrsf | 0.38 | 0.53 | **0.60** | 0.65 | 0.77 | 0.75 | 1 |
| LUAD-hofree | 0.34 | 0.50 | 0.56 | 0.62 | 0.78 | 1 | 1 |

**Table 3:** The comparison of RFSs models' performances that are trained with different feature representations for different cancer types. The methods are compared at the mean with four different quartile values of the 100 repeated runs, where each run is conducted with a random split of train and test data. The p-values of the one-tailed Wilcoxon signed-rank test of comparing the each model with Individual model is listed. The BH column lists the adjusted p-value after Benjamini and Hochberg correction for multiple hypothesis test correction [Benjamini and Hochberg(1995)].

#### 1.3 Performance Comparison of proposed PRER representation with Ternary PRER

| Cancer Type | p-value (compared to PRER) | BH adjusted p-value (compared to PRER) |
|---|---|---|
| **BRCA** | 0.9 | 1 |
| **BLCA** | 1 | 1 |
| **COAD** | 0.07 | 0.23 |
| **HNSC** | 0.21 | 0.41 |
| **GBM** | 0.01 | 0.08 |
| **KIRC** | 0.99 | 1 |
| **LUAD** | 0.80 | 1 |
| **LUSC** | 0.99 | 1 |
| **OV** | 0.11 | 0.27 |
| **UCEC** | 0.02 | 0.08 |

**Table 4:** Comparison of RSF model performances that are trained with two PRER settings for each cancer type. The performance of the proposed PRER repsesentation (PRER) is compared with the PRER representation with three value as features (Ternary PRER) in each case. Wilcoxon signed-rank test is applied to calculate p-value for of each setting with respect to PRER representation. The BH column lists the adjusted p-value after Benjamini and Hochberg correction for multiple hypothesis test correction [Benjamini and Hochberg (1995)].

#### 1.4 Prediction Performance with IntAct PPI Network

| **Random Survival Forest C-indices** | | | | | | | |
|---|---|---|---|---|---|---|---|
| | C-index | | | | | p-value | BH adjusted p-value |
| | $1^{st}$ | $2^{nd}$ | median | $3^{rd}$ | $4^{th}$ | | |
| KIRC | 0.57 | 0.66 | 0.69 | 0.72 | 0.80 | **$1.1\text{x}10^{-8}$** | **$3.6\text{x}10^{-8}$** |
| KIRC-prer | 0.56 | 0.68 | **0.70** | 0.73 | 0.83 | | |
| OV | 0.40 | 0.52 | 0.55 | 0.57 | 0.66 | **0.21** | 0.30 |
| OV-prer | 0.44 | 0.52 | 0.55 | 0.58 | 0.66 | | |
| BRCA | 0.38 | 0.55 | 0.60 | 0.64 | 0.82 | **$8.3\text{x}10^{-8}$** | **$1.7\text{x}10^{-7}$** |
| BRCA-prer | 0.47 | 0.58 | **0.64** | 0.70 | 0.80 | | |
| LUSC | 0.36 | 0.51 | 0.57 | 0.64 | 0.73 | **$5.3\text{x}10^{-9}$** | **$2.7\text{x}10^{-8}$** |
| LUSC-prer | 0.44 | 0.58 | **0.63** | 0.68 | 0.78 | | |
| COAD | 0.22 | 0.43 | 0.49 | 0.53 | 0.70 | **$1.2\text{x}10^{-4}$** | **$2.0\text{x}10^{-4}$** |
| COAD-prer | 0.34 | 0.46 | **0.52** | 0.57 | 0.70 | | |
| HNSC | 0.37 | 0.48 | 0.52 | 0.55 | 0.70 | **$3.9\text{x}10^{-8}$** | **$9.9\text{x}10^{-8}$** |
| HNSC-prer | 0.38 | 0.52 | **0.56** | 0.60 | 0.71 | | |
| UCEC | 0.27 | 0.47 | 0.53 | 0.59 | 0.80 | **$3.6\text{x}10^{-17}$** | **$3.6\text{x}10^{-16}$** |
| UCEC-prer | 0.46 | 0.61 | **0.66** | 0.70 | 0.86 | | |
| GBM | 0.35 | 0.51 | 0.56 | 0.61 | 0.70 | **0.47** | 0.59 |
| GBM-prer | 0.43 | 0.53 | **0.57** | 0.62 | 0.71 | | |
| BLCA | 0.31 | 0.47 | 0.55 | 0.61 | 0.83 | **0.99** | 1 |
| BLCA-prer | 0.27 | 0.45 | **0.51** | 0.57 | 0.77 | | |
| LUAD | 0.40 | 0.55 | 0.59 | 0.66 | 0.80 | 1 | 1 |
| LUAD-prer | 0.23 | 0.42 | 0.48 | 0.54 | 0.81 | | |

**Table 5:** The comparison of RFSs models' performances that are trained with individual features and pairwise ranking representations (PRER) in IntAct PPI for different cancer types. The methods are compared at the mean with four different quartile values of the 100 repeated runs, where each run is conducted with a random split of train and test data. The p-values of the one-tailed Wilcoxon signed-rank test of comparing the two models is listed. The BH column lists the adjusted p-value after Benjamini and Hochberg correction for multiple hypothesis test correction[Benjamini and Hochberg(1995)].

### 2 Supplementary Figures

#### 2.1 PRER Compared to Alternative Methods

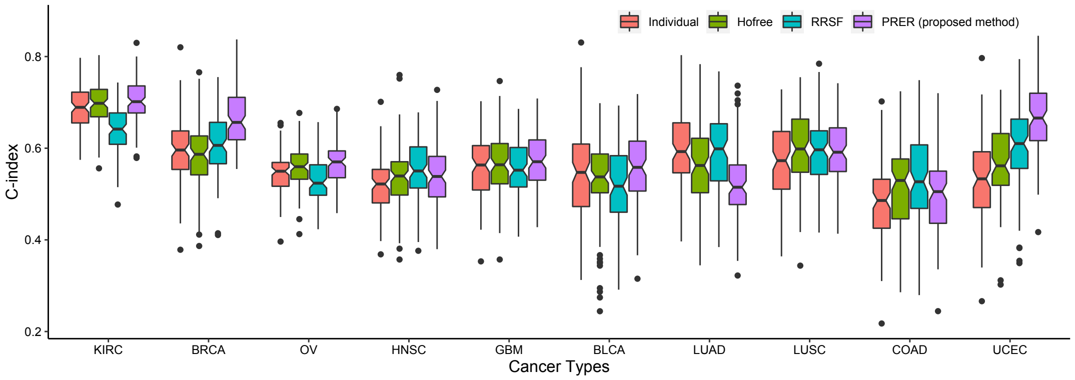


**Figure 1:** Comparison of different RSF prediction models. Hofree: a model that uses network propagated protein expressions as features, Individual: a model that is trained with individual protein expressions, PRER: the proposed PRER representation, RRSF: a model that is proposed by Wang et al. 2018.

#### 2.2 Performance Comparisons of Two PRER Settings

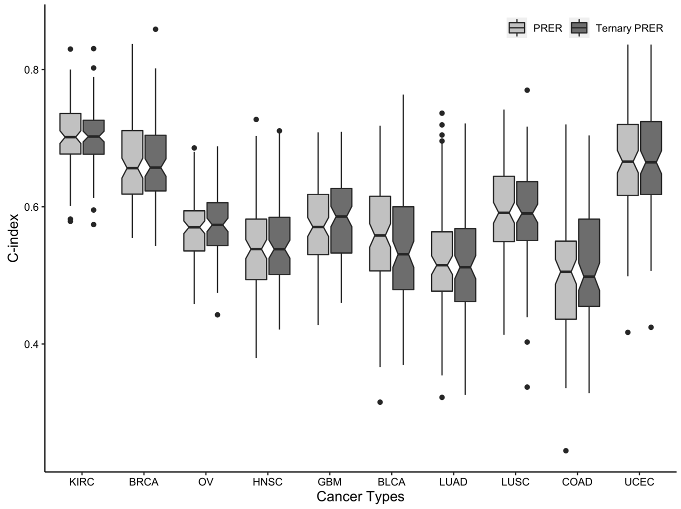


**Figure 2:** Comparison of different models trained with different PRER settings for each cancer type. PRER: the proposed PRER representation (PRER), Ternary PRER: PRER representation with three different assignment of feature values (-1, 0, 1). The box-plots shows the distribution over 100 models that are trained on different random train/test splits. The models are compared statistically in Supplementary Table 4.

#### 2.3 The Effect of Random Walk Parameters on Performance

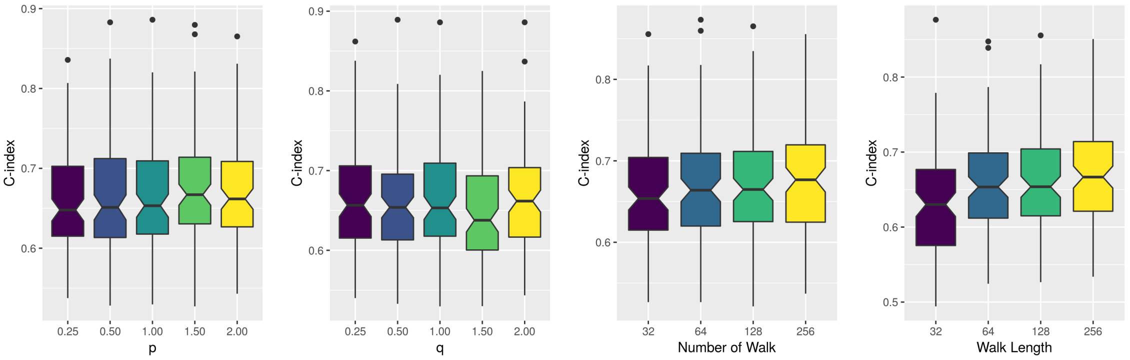


(a) (b) (c) (d)

**Figure 3:** Parameter sensitivity for BRCA. At each subfigure, corresponding parameter is varied and all other parameters are fixed.

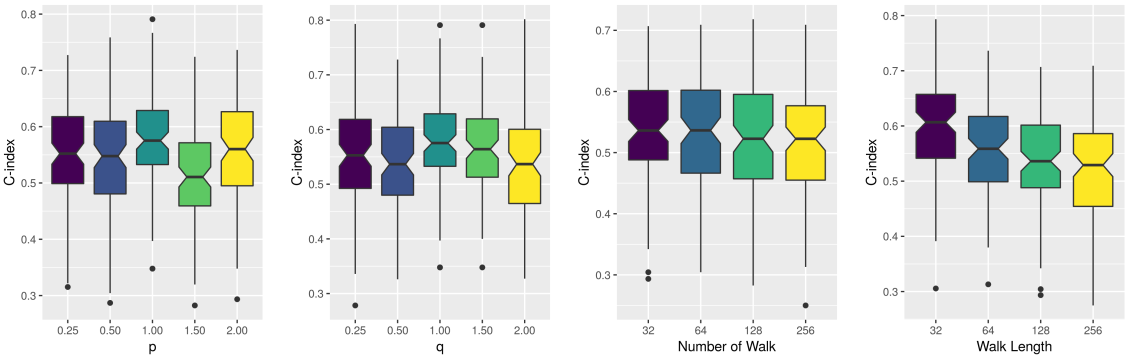


(a) (b) (c) (d)

**Figure 4:** Parameter sensitivity for BLCA. At each subfigure, corresponding parameter is varied and all other parameters are fixed.

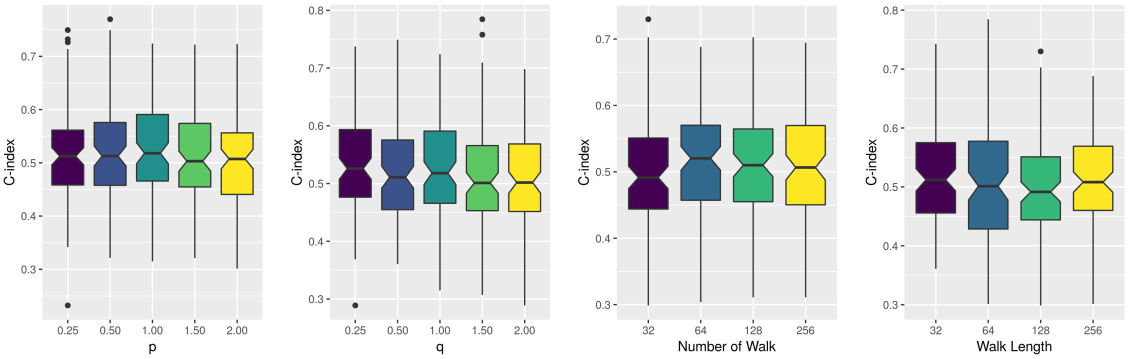


(a) (b) (c) (d)

**Figure 5:** Parameter sensitivity for COAD. At each subfigure, corresponding parameter is varied and all other parameters are fixed.

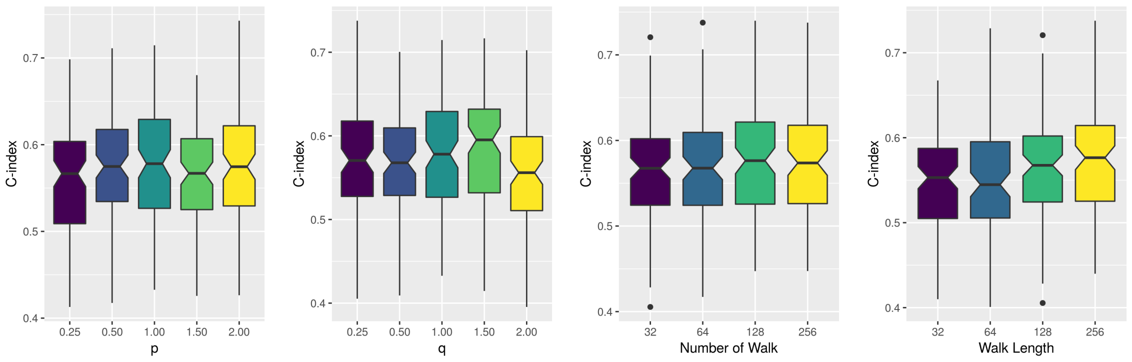


(a) (b) (c) (d)

**Figure 6:** Parameter sensitivity for GBM. At each subfigure, corresponding parameter is varied and all other parameters are fixed.

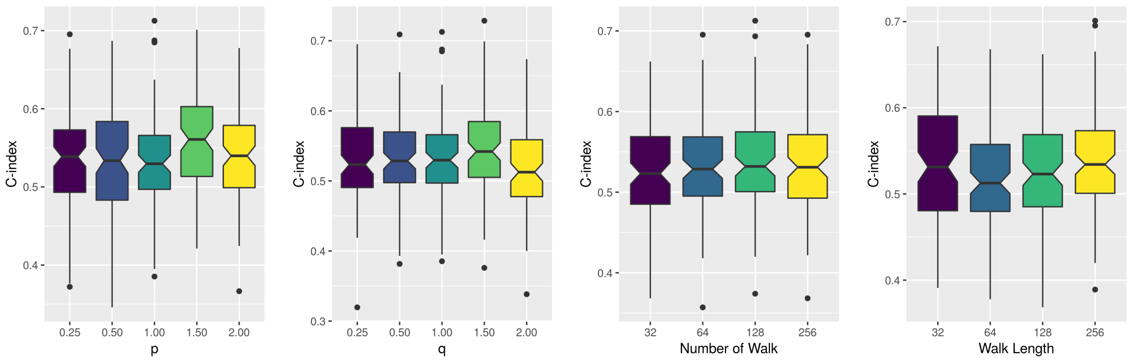


(a) (b) (c) (d)

**Figure 7:** Parameter sensitivity for HNSC. At each subfigure, corresponding parameter is varied and all other parameters are fixed.

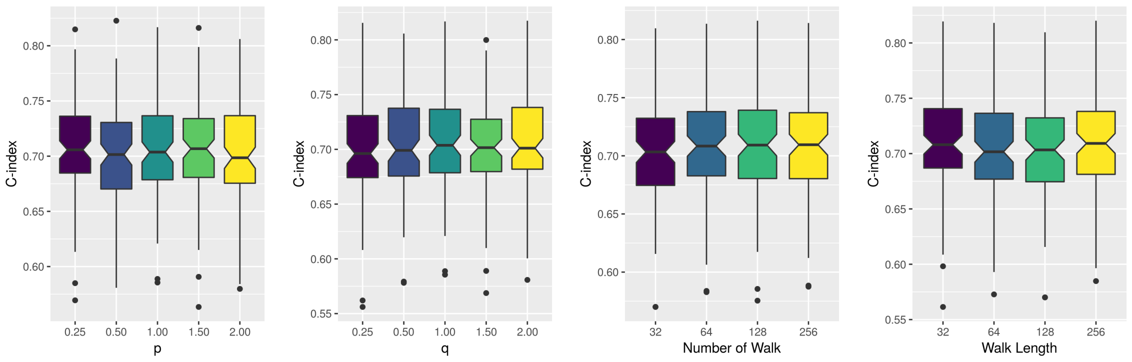


(a) (b) (c) (d)

**Figure 8:** Parameter sensitivity for KIRC. At each subfigure, corresponding parameter is varied and all other parameters are fixed.

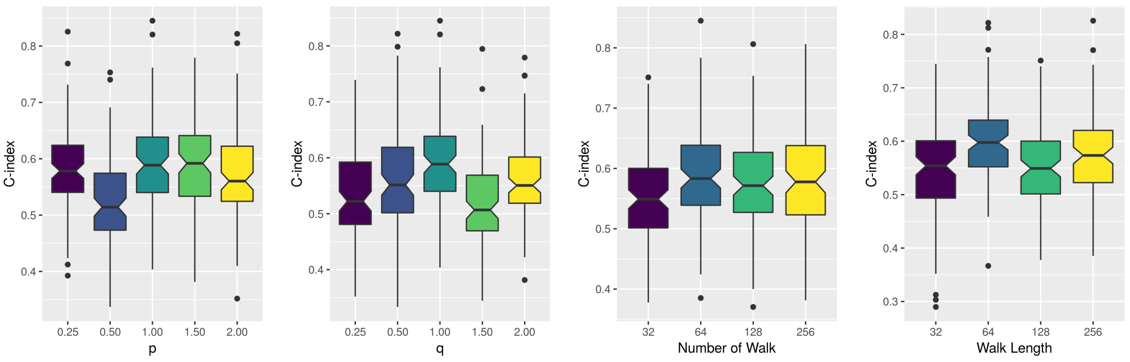


(a) (b) (c) (d)

**Figure 9:** Parameter sensitivity for LUAD. At each subfigure, corresponding parameter is varied and all other parameters are fixed.

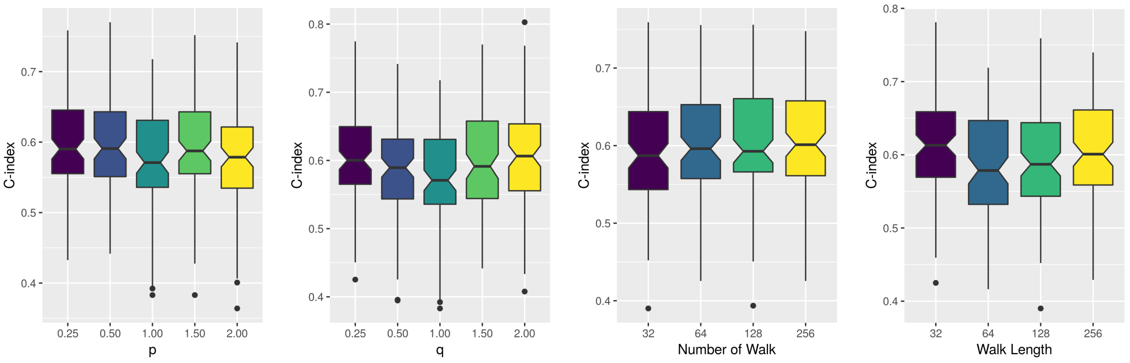


(a) (b) (c) (d)

**Figure 10:** Parameter sensitivity for LUSC. At each subfigure, corresponding parameter is varied and all other parameters are fixed.

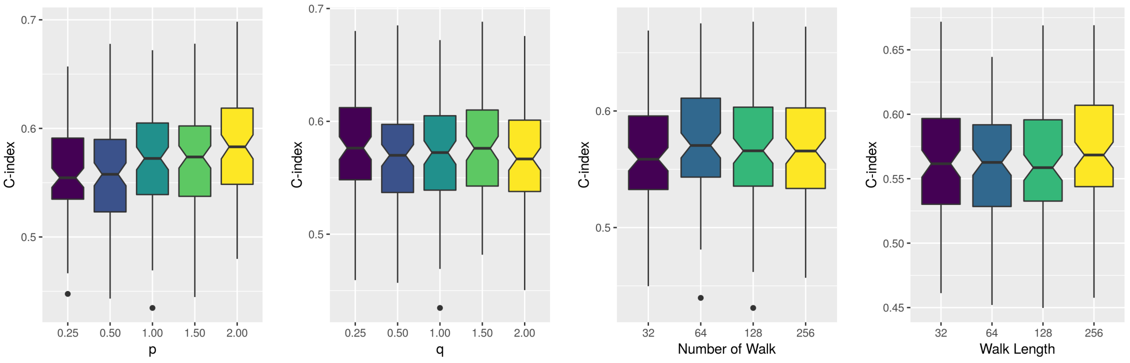


(a) (b) (c) (d)

**Figure 11:** Parameter sensitivity for OV. At each subfigure, corresponding parameter is varied and all other parameters are fixed.

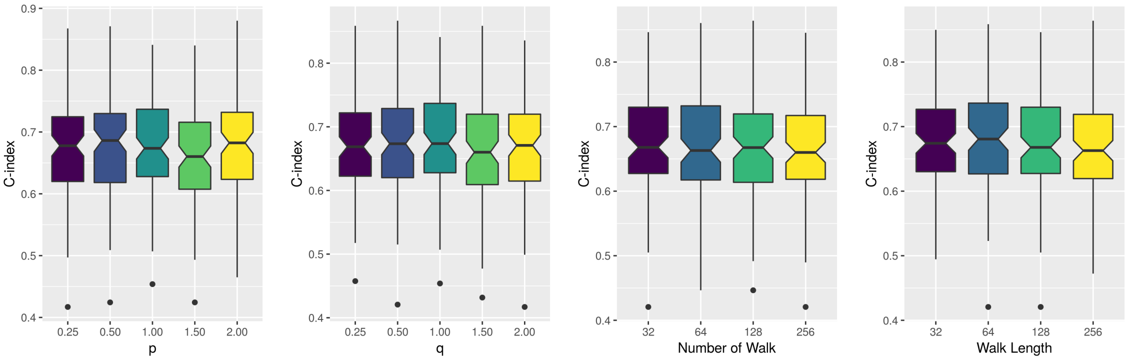


(a) (b) (c) (d)

**Figure 12:** Parameter sensitivity for UCEC. At each subfigure, corresponding parameter is varied and all other parameters are fixed.

#### 2.4 Important PRER Features

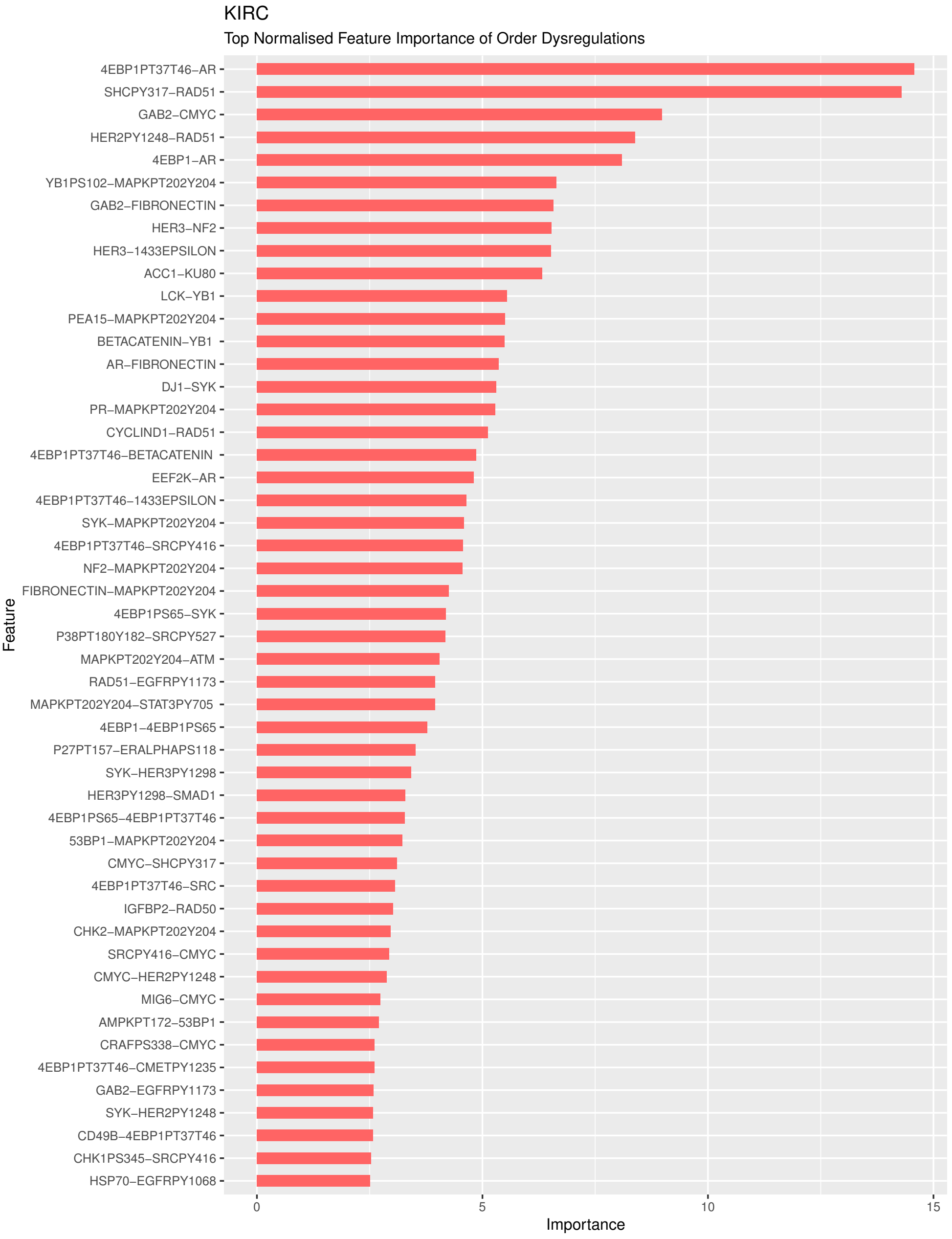


**Figure 13:** Variable importance of significant pairwise ranking embeddings for kidney renal clear cell carcinoma.

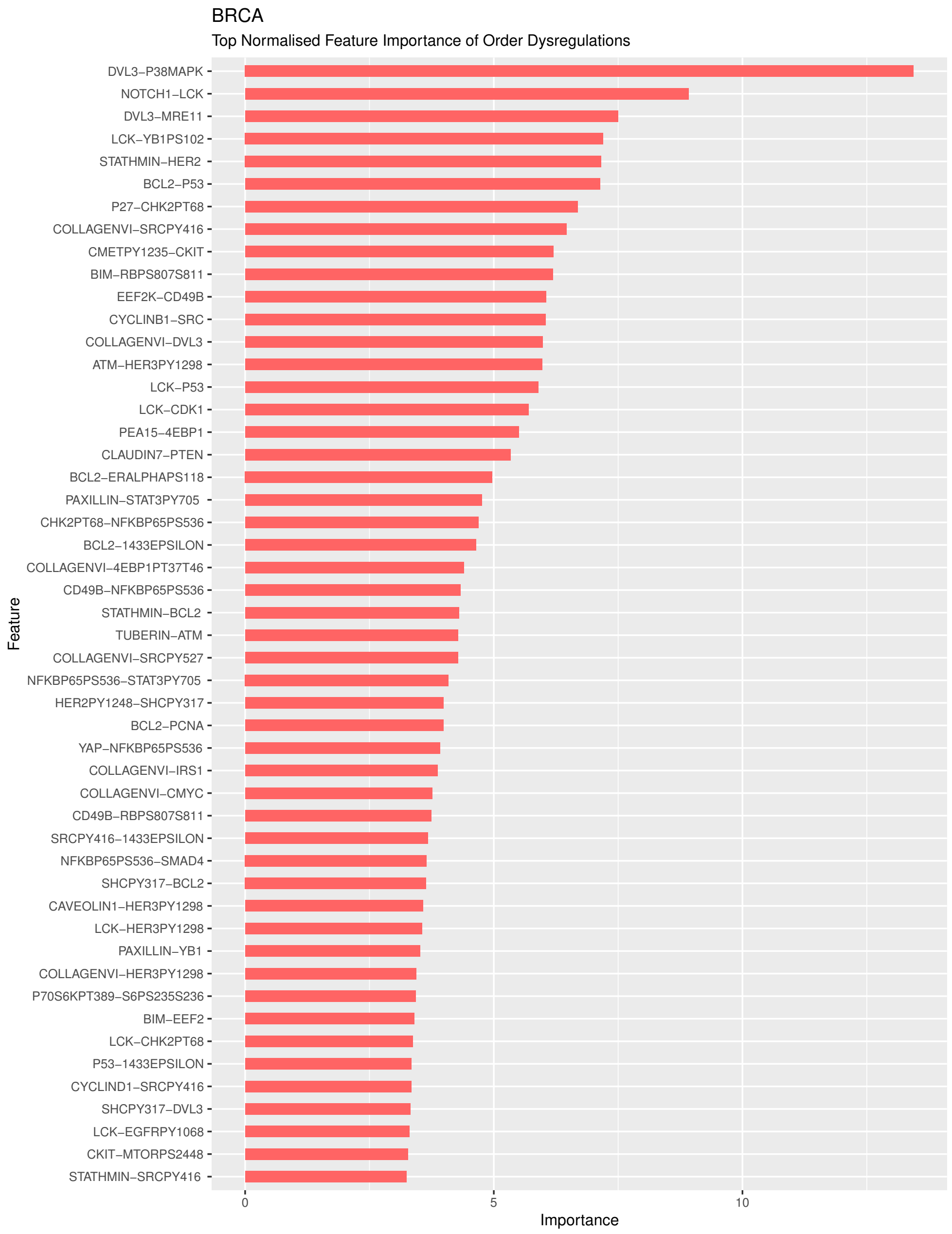


**Figure 14:** Variable importance of significant pairwise ranking embeddings for breast invasive carcinoma.

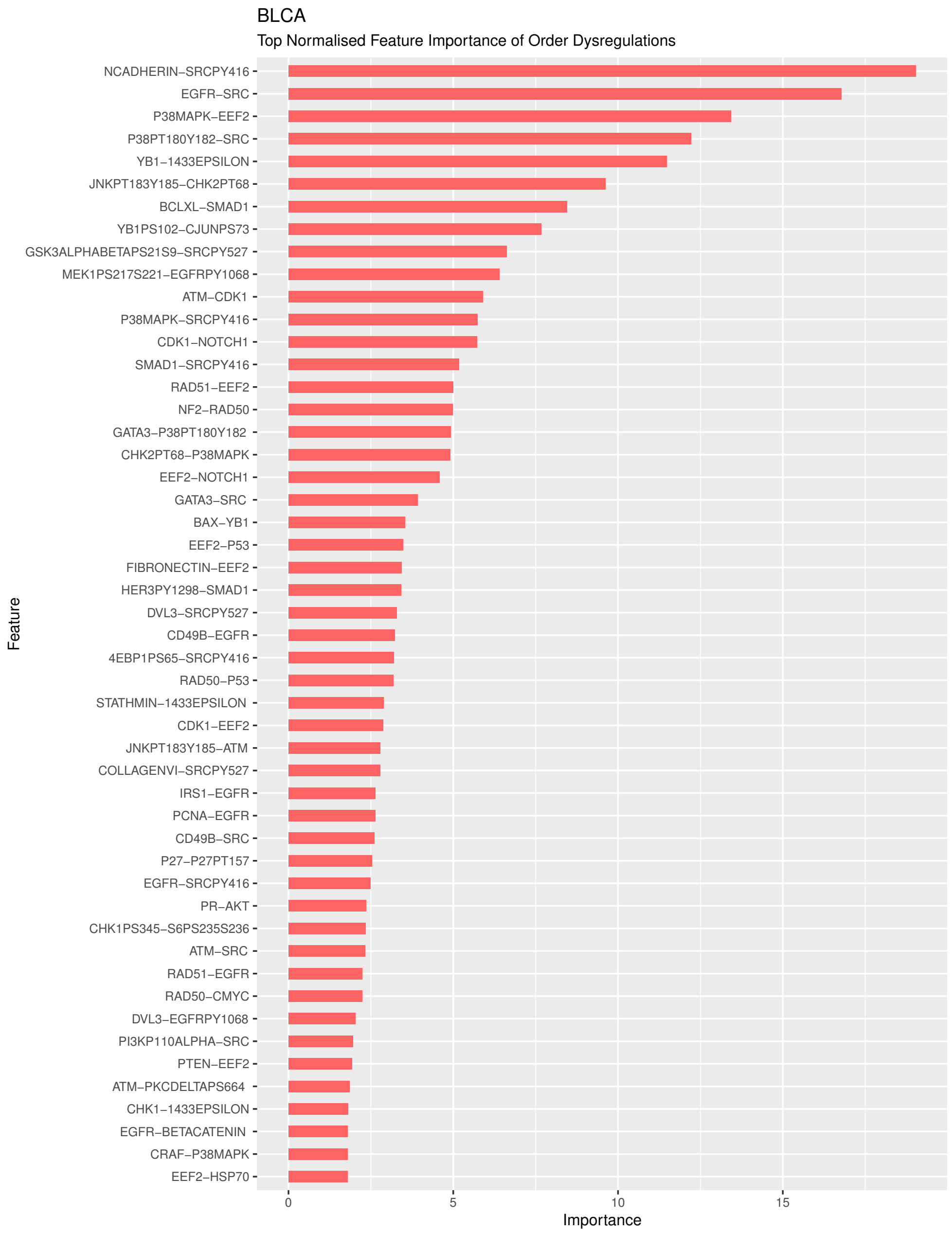


**Figure 15:** Variable importance of significant pairwise ranking embeddings for bladder urothelial carcinoma.

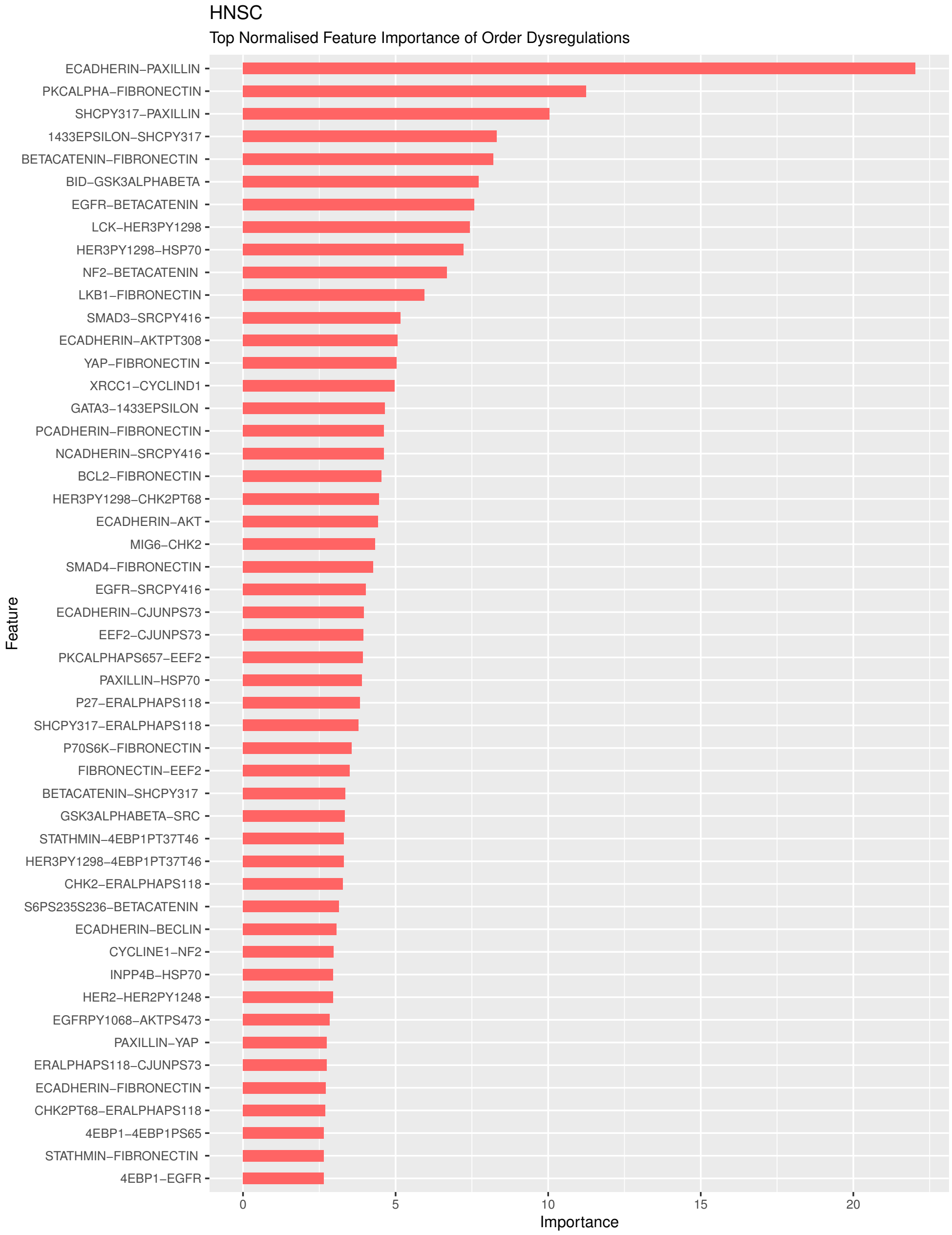


**Figure 16:** Variable importance of significant pairwise ranking embeddings for head and neck squamous cell carcinoma.

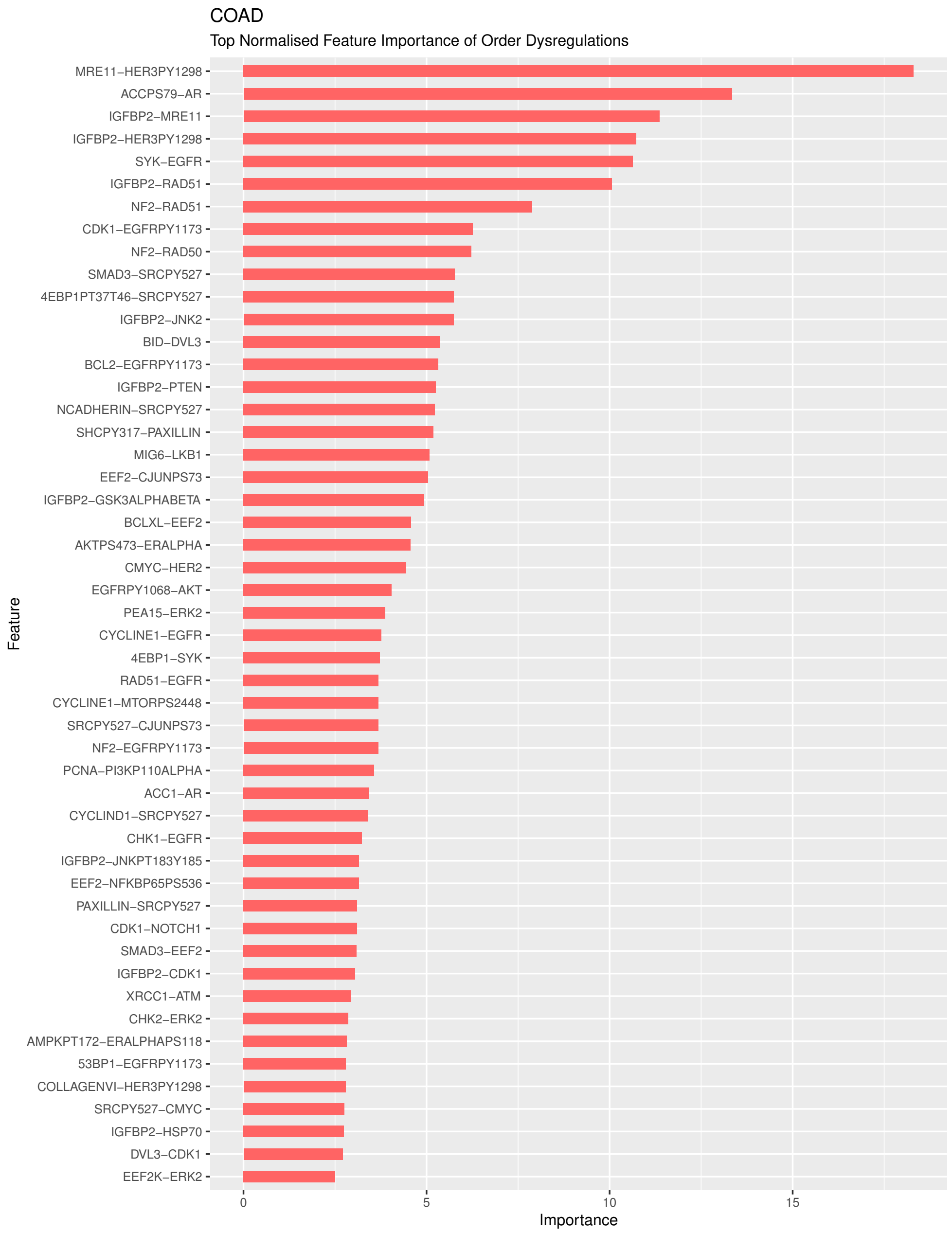


**Figure 17:** Variable importance of significant pairwise ranking embeddings for colon adenocarcinoma.

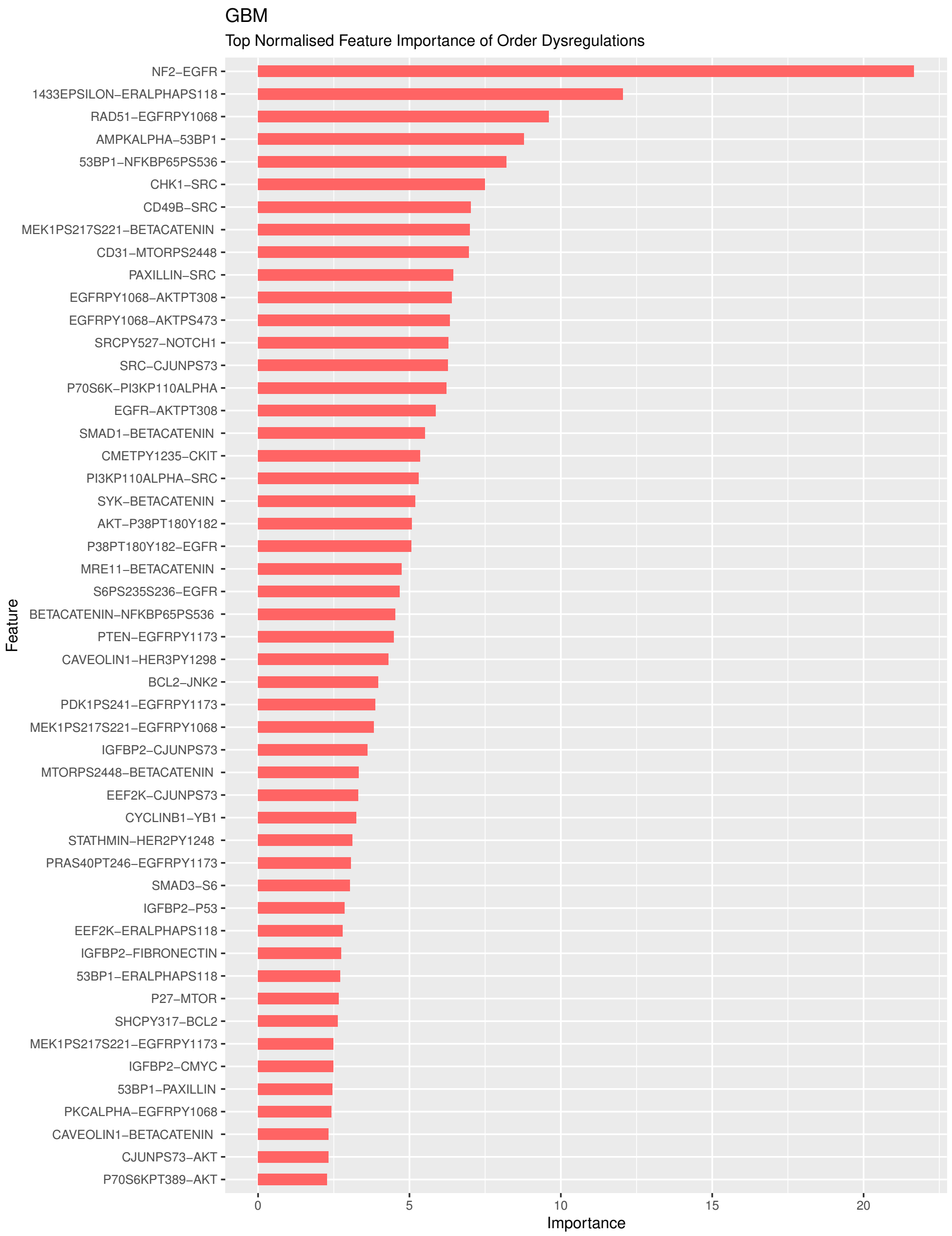


**Figure 18:** Variable importance of significant pairwise ranking embeddings for glioblastoma multiforme.

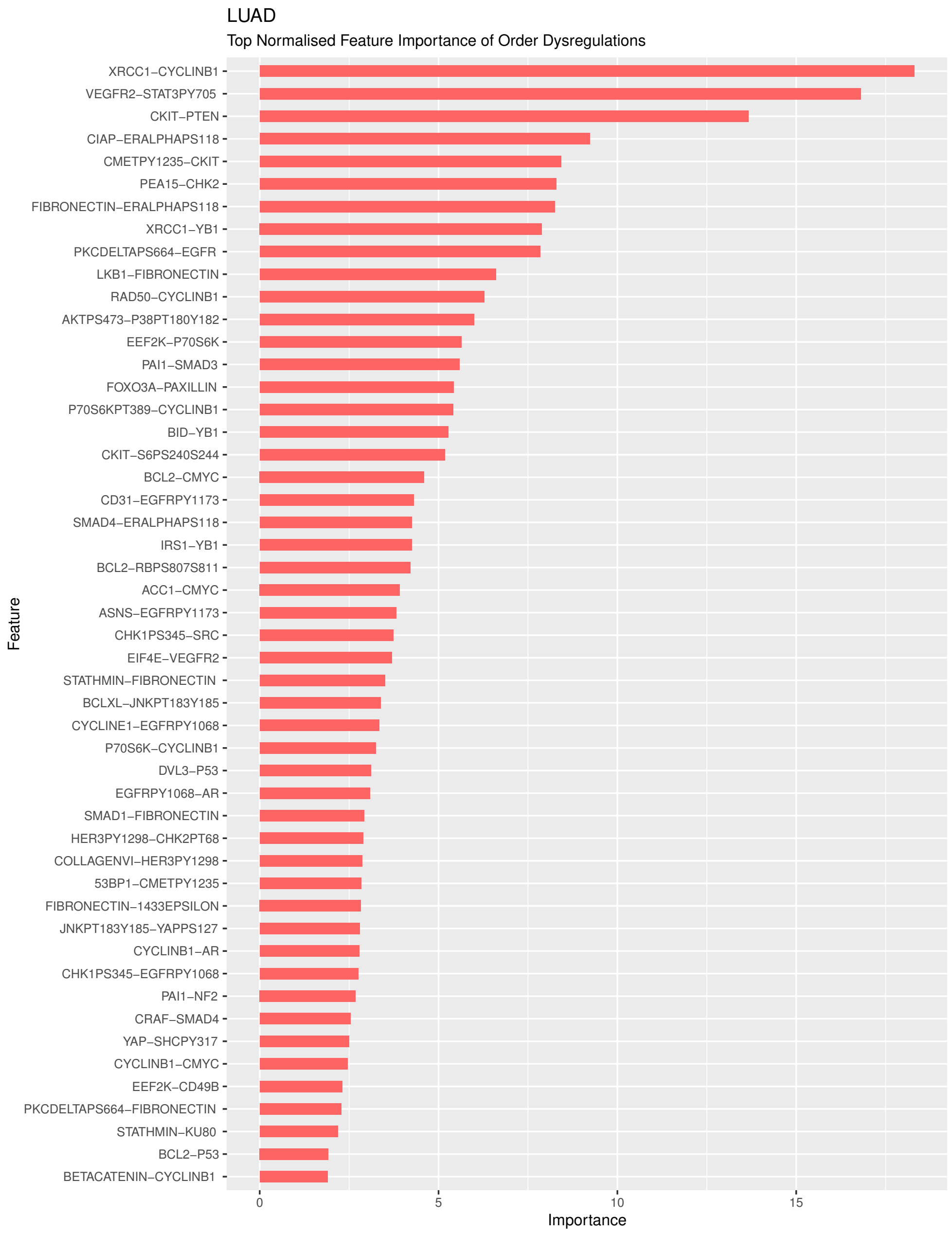


**Figure 19:** Variable importance of significant pairwise ranking embeddings for lung adenocarcinoma.

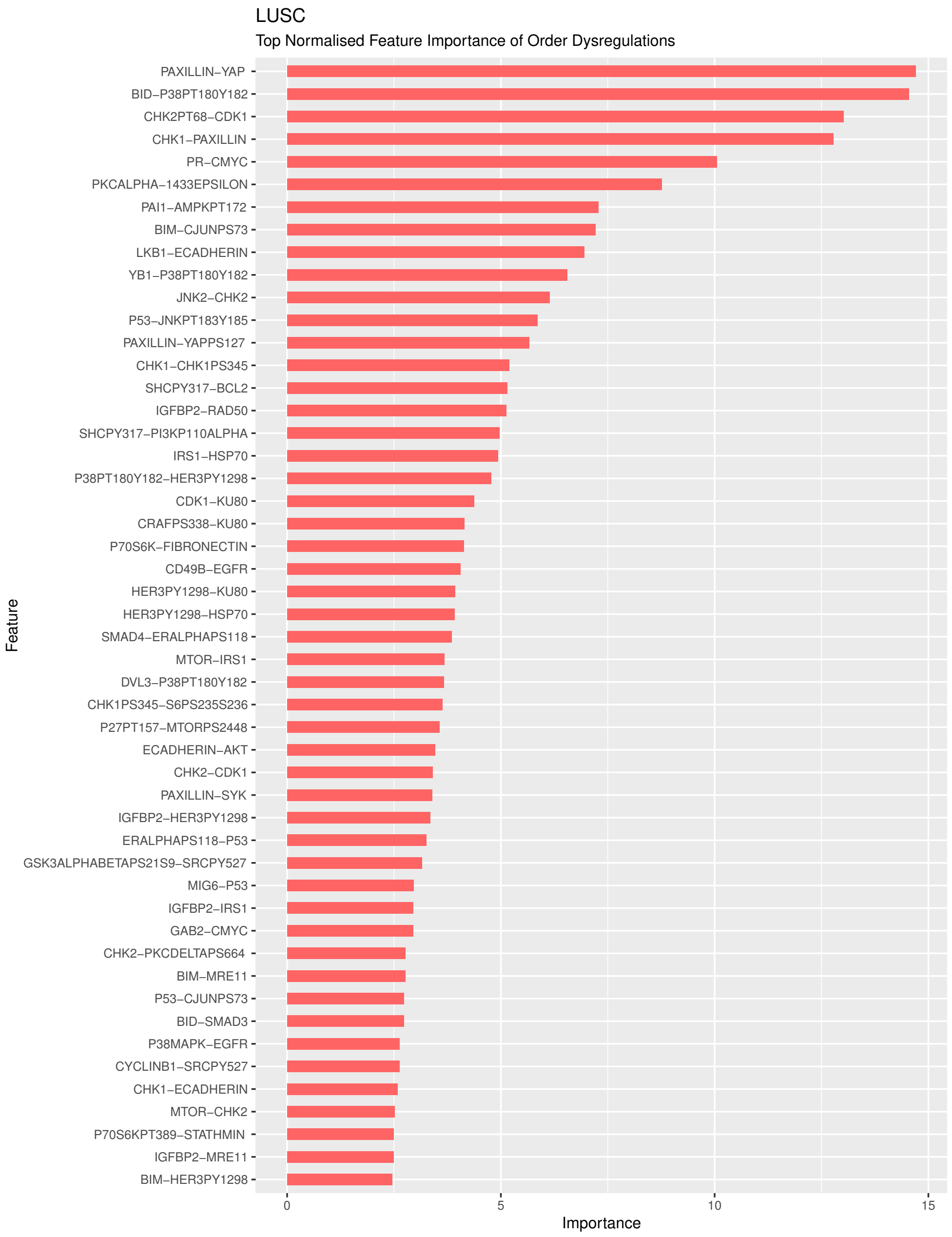


**Figure 20:** Variable importance of significant pairwise ranking embeddings for lung squamous cell carcinoma.

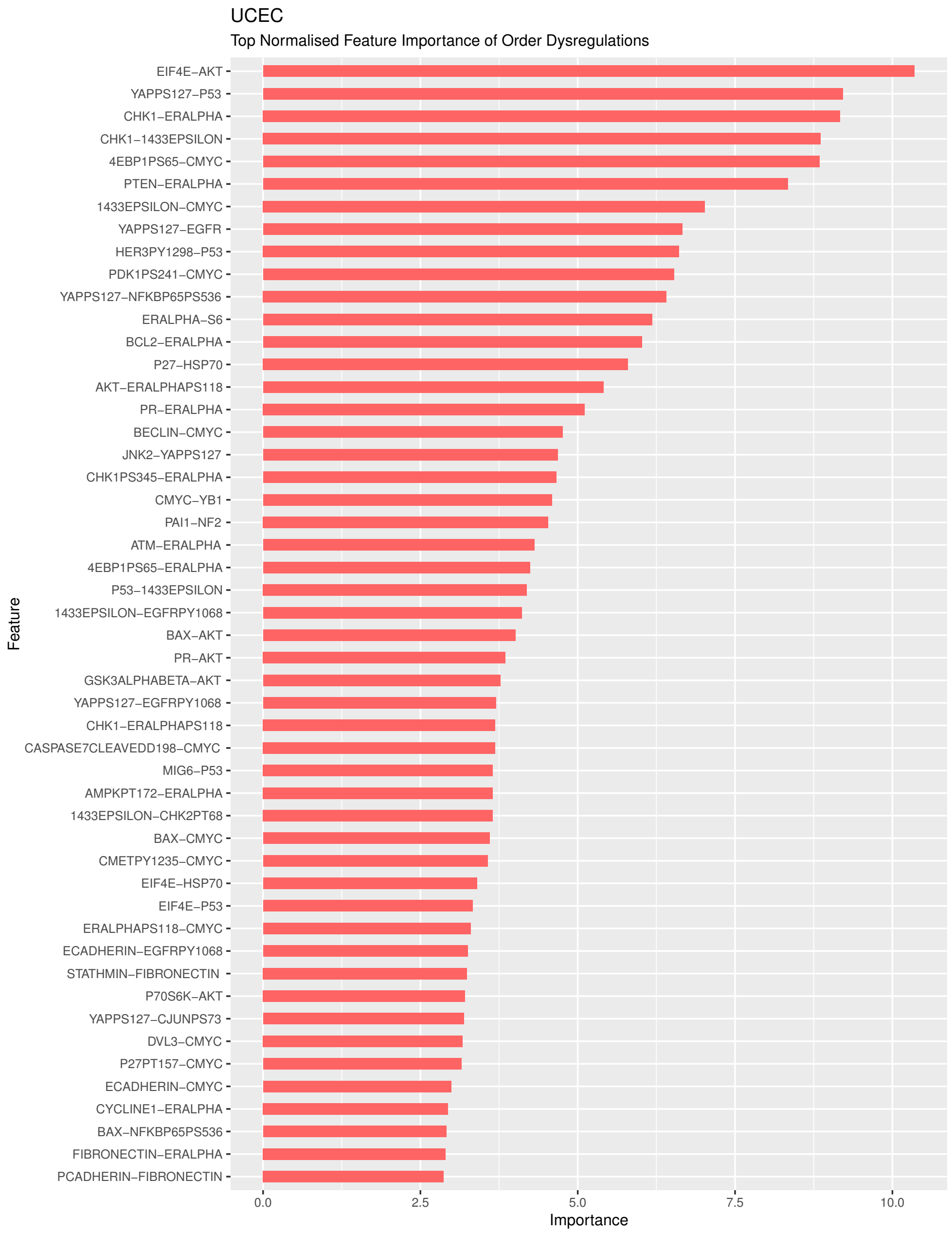


**Figure 21:** Variable importance of significant pairwise ranking embeddings for uterine corpus endometrial carcinoma.

#### 2.5 PRER Networks

##### KIRC

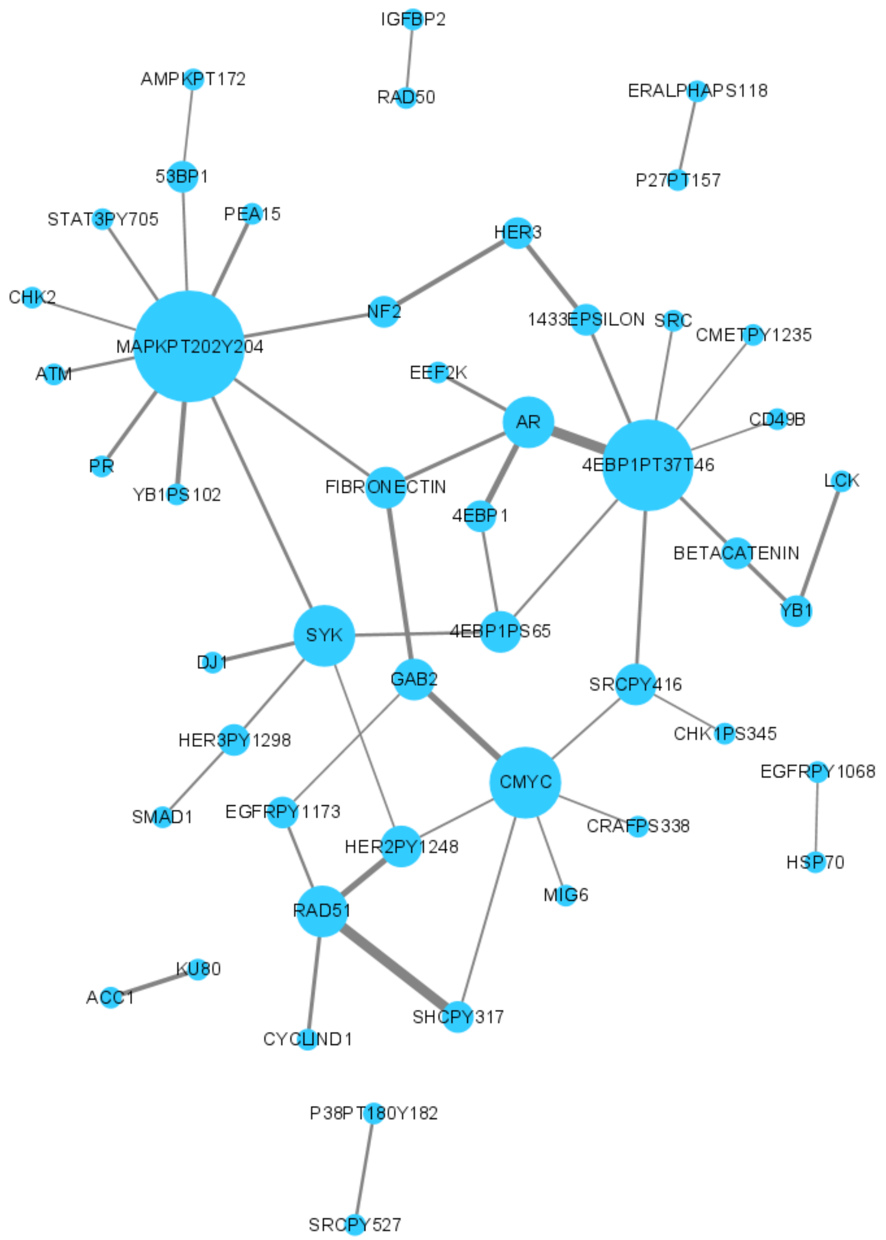


**Figure 22:** Nodes represent proteins that appear in the top 50 pairwise ranking embeddings for kidney renal clear cell carcinoma; edges represent that two proteins participate in a pairwise rank order feature together.

**BRCA**

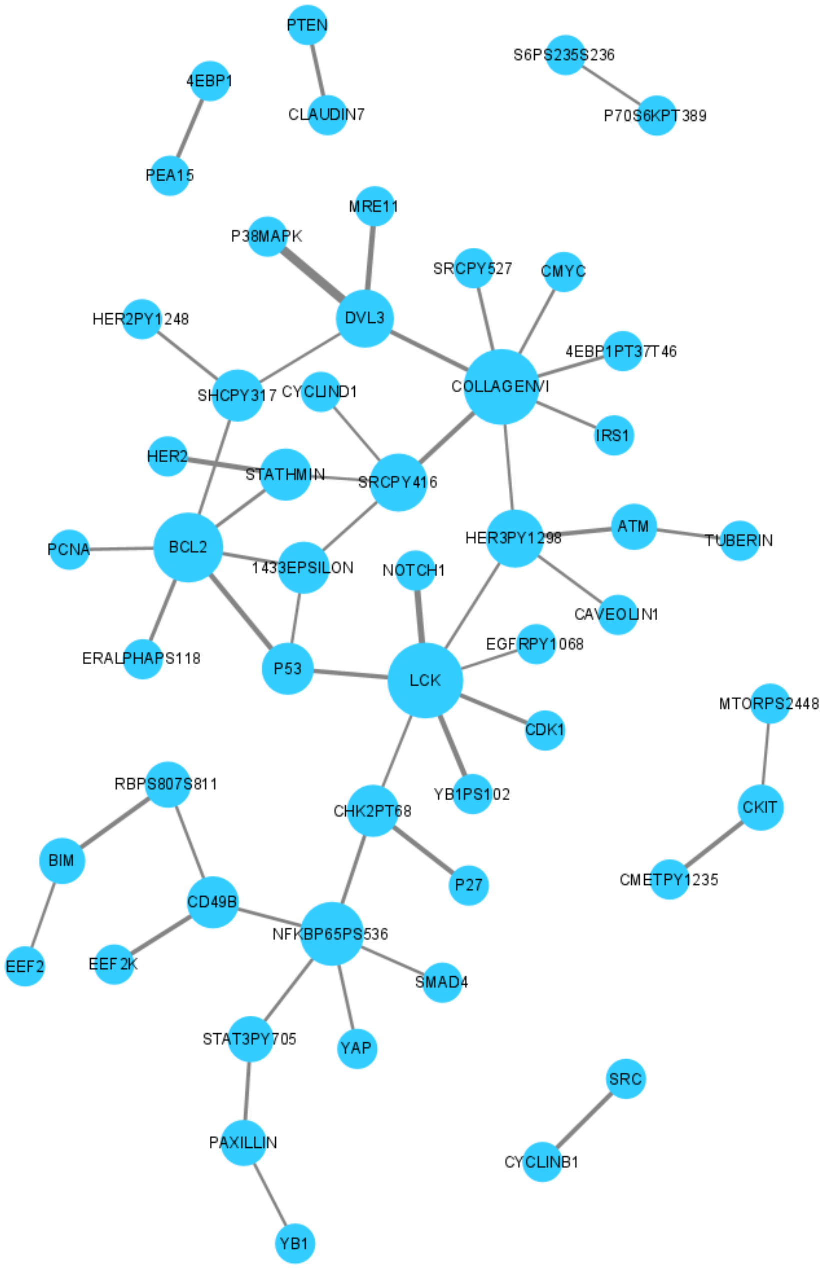


**Figure 23:** Nodes represent proteins that appear in the top 50 pairwise ranking embeddings for breast invasive carcinoma; edges represent that two proteins participate in a pairwise rank order feature together.

**BLCA**

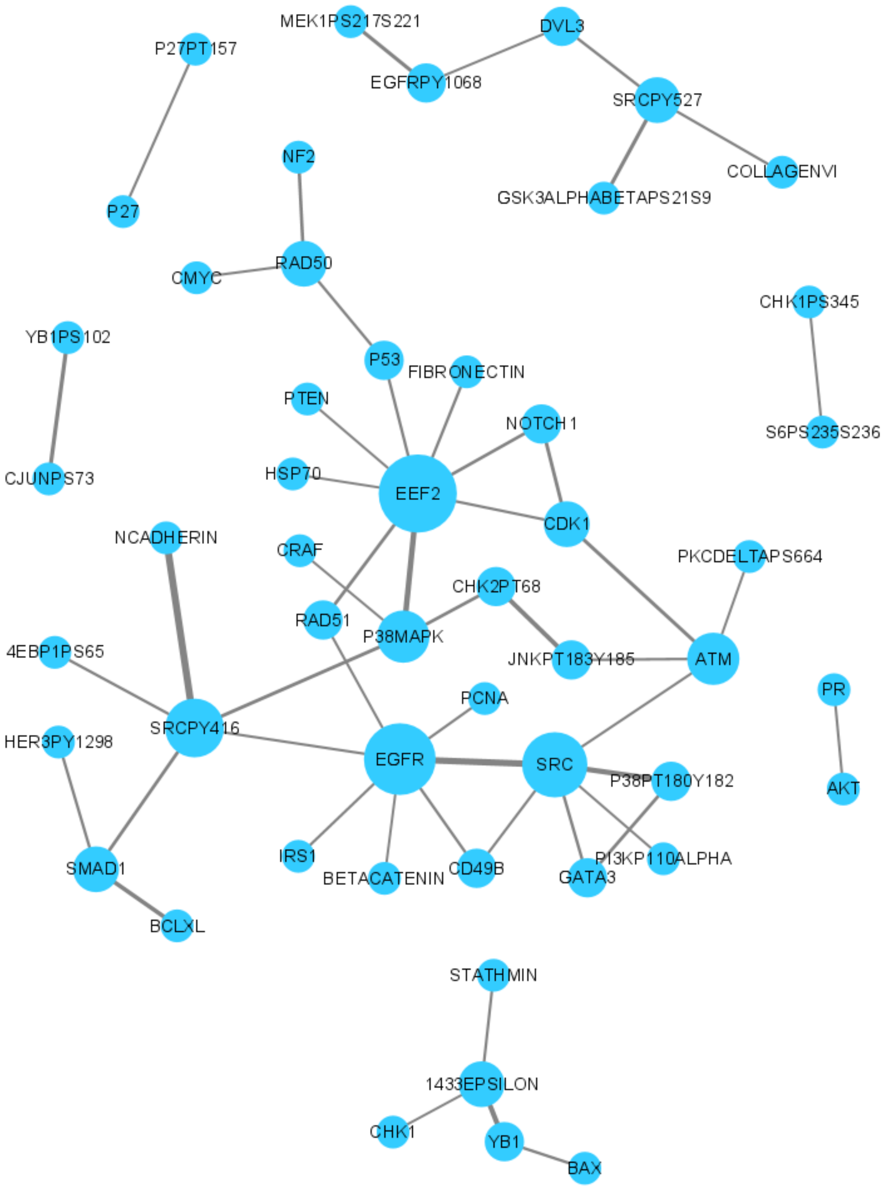


**Figure 24:** Nodes represent proteins that appear in the top 50 pairwise ranking embeddings for bladder urothelial carcinoma; edges represent that two proteins participate in a pairwise rank order feature together.

**HNSC**

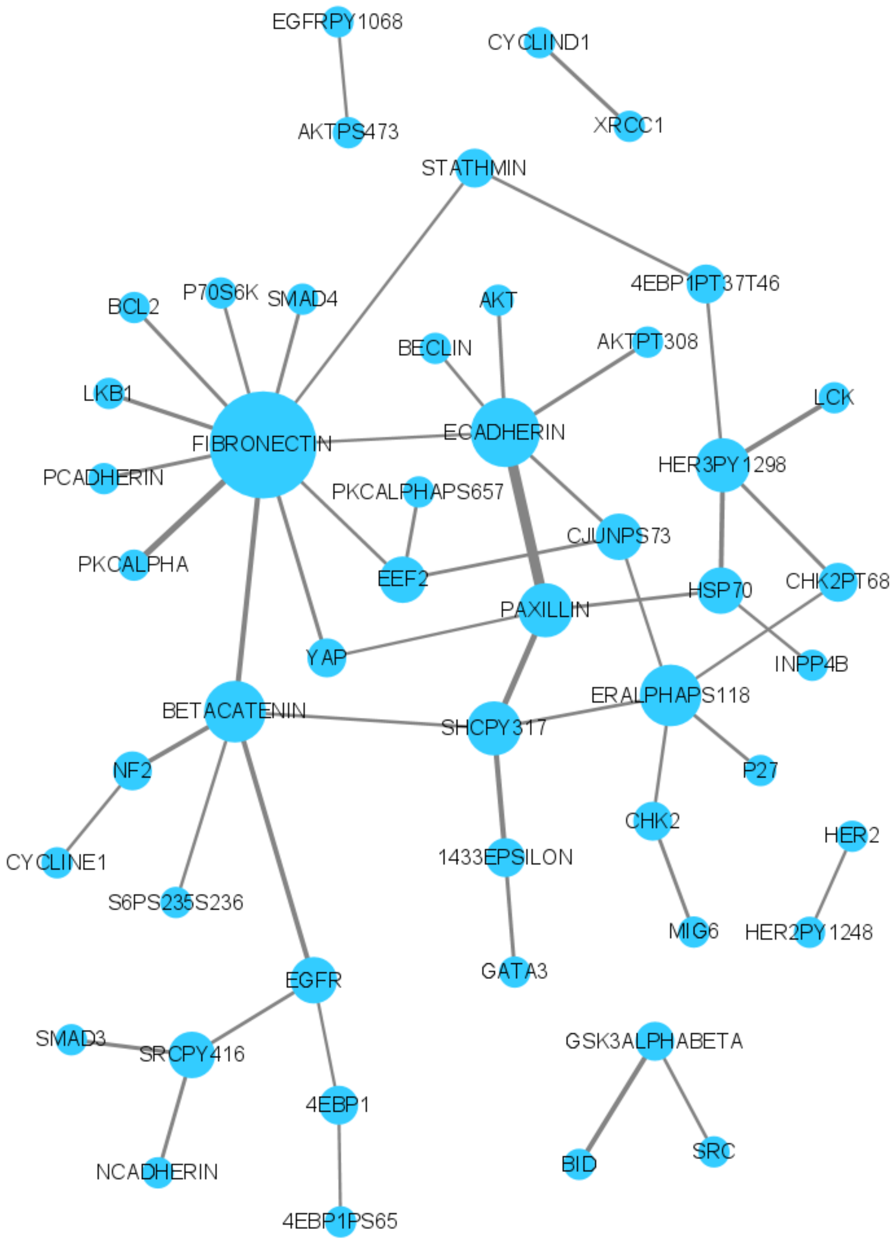


**Figure 25:** Nodes represent proteins that appear in the top 50 pairwise ranking embeddings for head and neck squamous cell carcinoma; edges represent that two proteins participate in a pairwise rank order feature together.

##### COAD

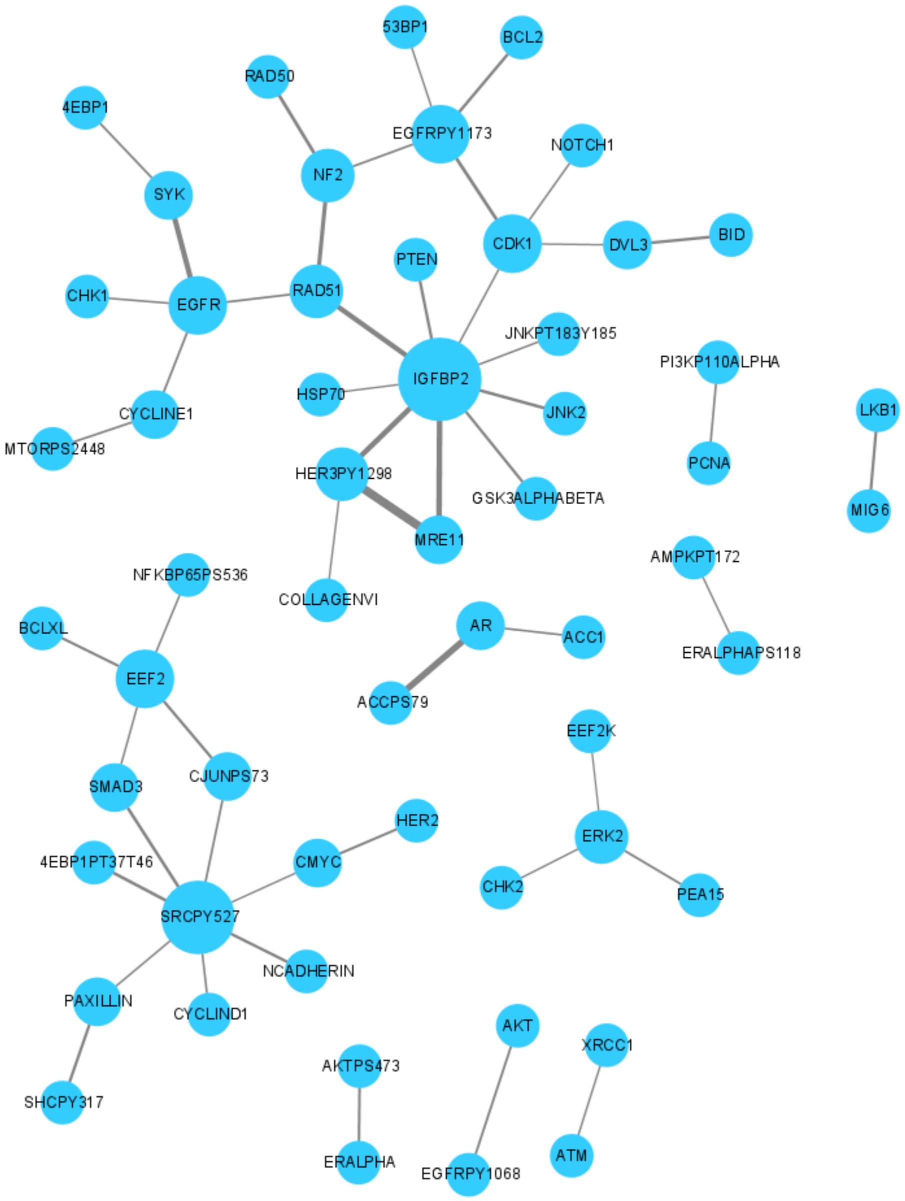


**Figure 26:** Nodes represent proteins that appear in the top 50 pairwise ranking embeddings for colon adenocarcinoma; edges represent that two proteins participate in a pairwise rank order feature together.

#### GBM

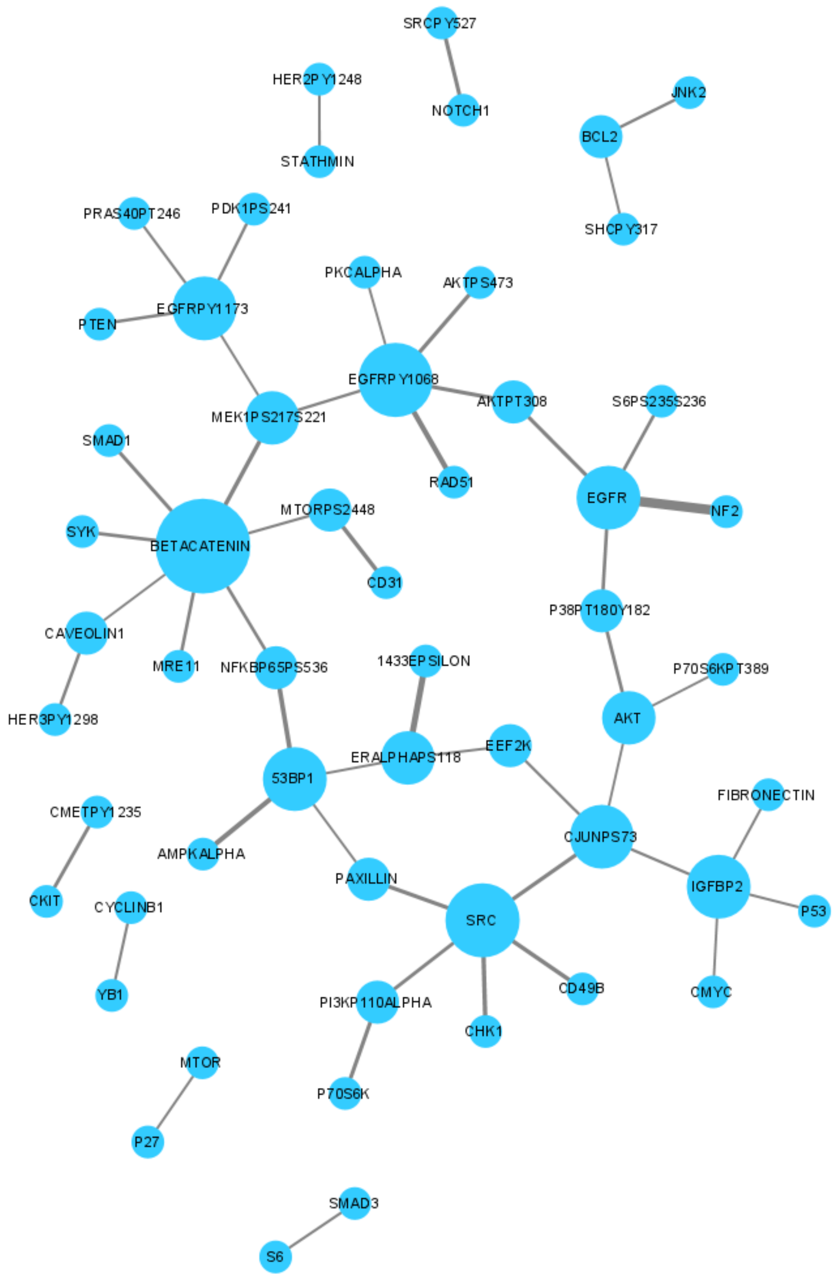


**Figure 27:** Nodes represent proteins that appear in the top 50 pairwise ranking embeddings for glioblastoma multiforme; edges represent that two proteins participate in a pairwise rank order feature together.

**LUAD**

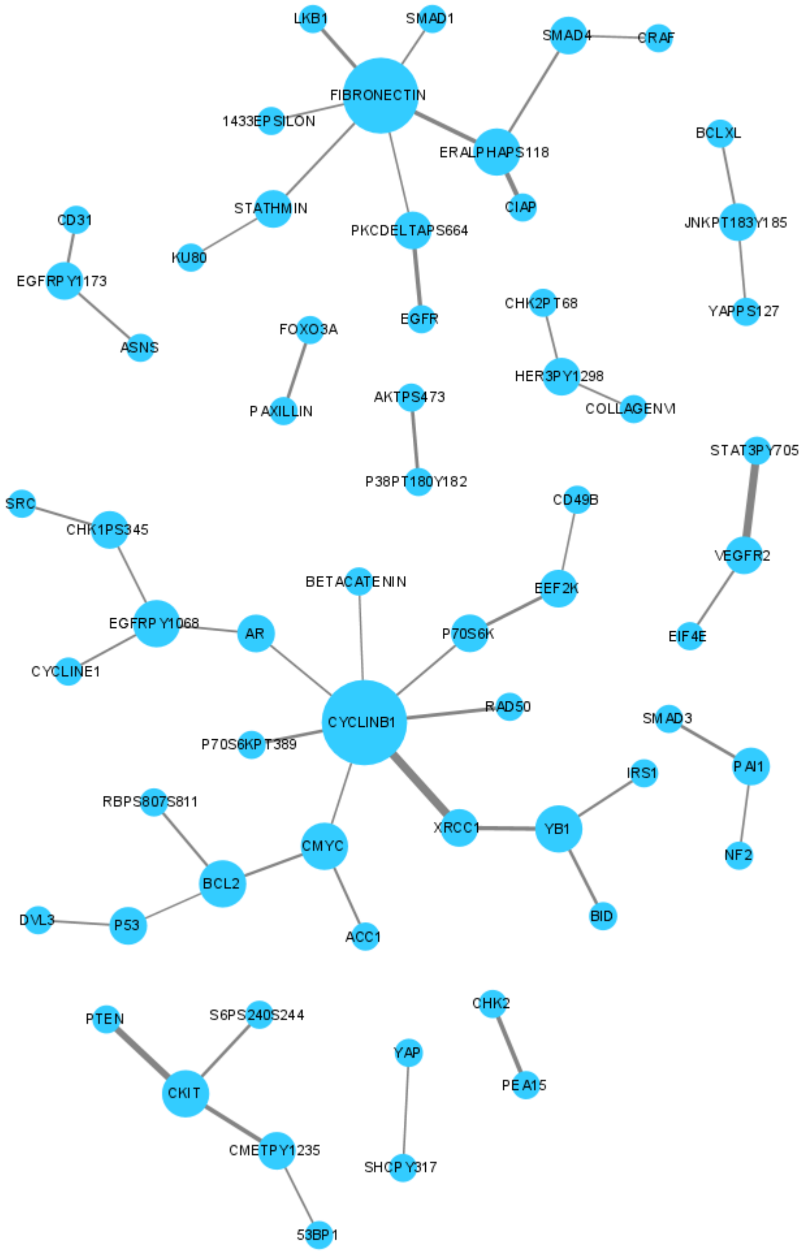


**Figure 28:** Nodes represent proteins that appear in the top 50 pairwise ranking embeddings for lung adenocarcinoma; edges represent that two proteins participate in a pairwise rank order feature together.

#### LUSC

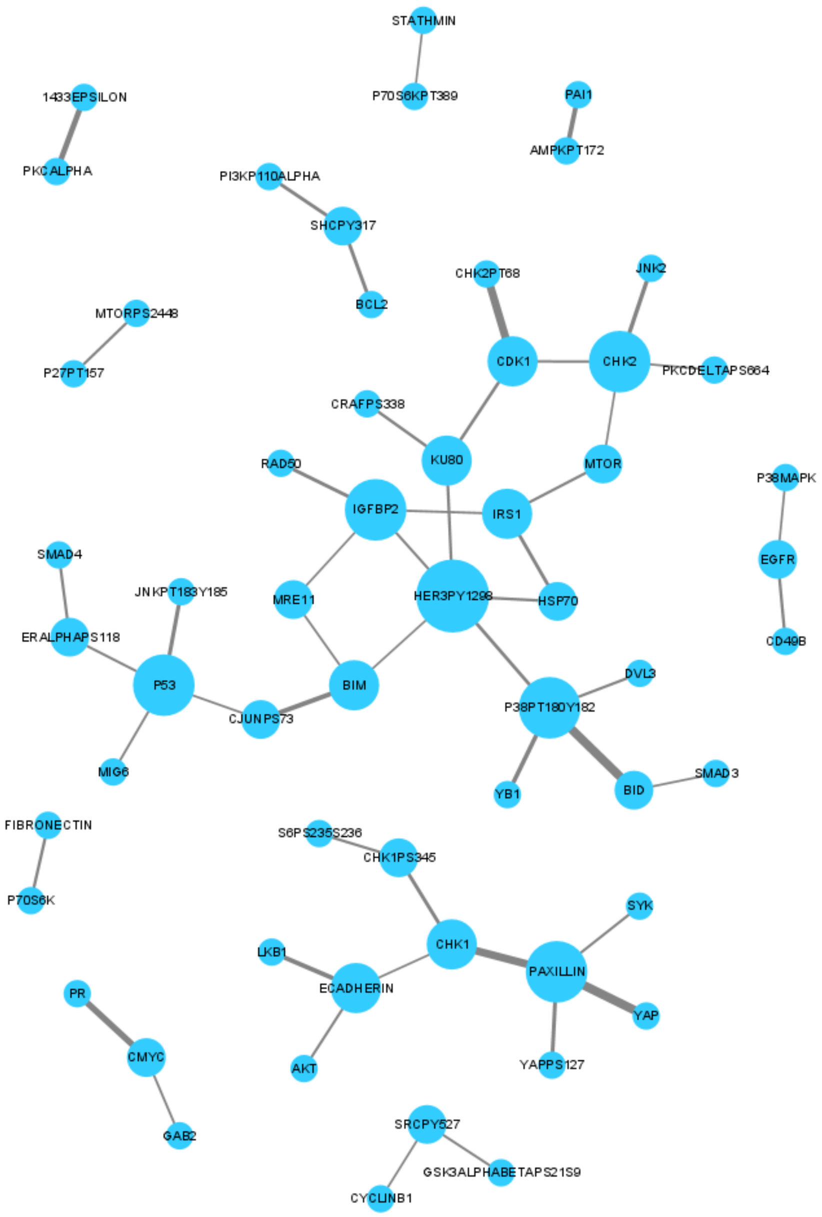


**Figure 29:** Nodes represent proteins that appear in the top 50 pairwise ranking embeddings for lung squamous cell carcinoma; edges represent that two proteins participate in a pairwise rank order feature together.

**UCEC**

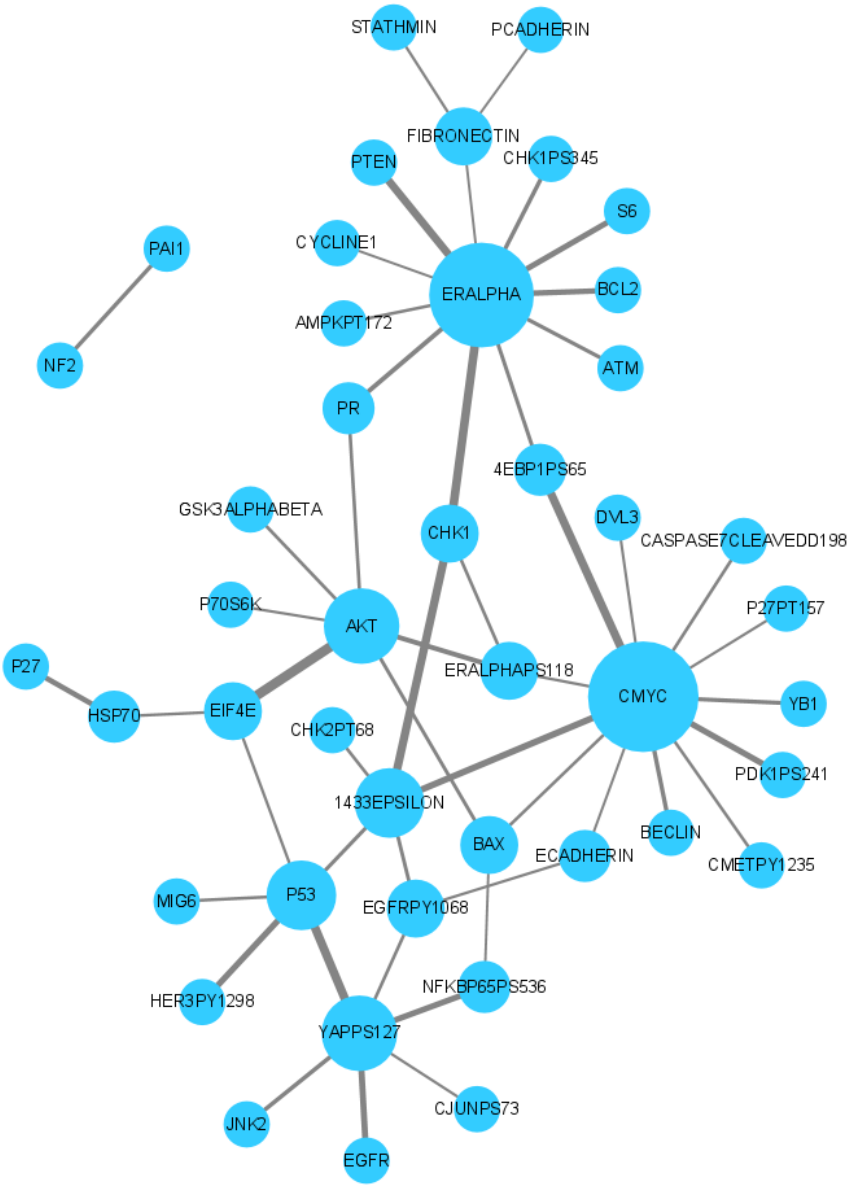


**Figure 30:** Nodes represent proteins that appear in the top 50 pairwise ranking embeddings for uterine corpus endometrial carcinoma; edges represent that two proteins participate in a pairwise rank order feature together.

#### 2.6 Top Predictive PRER are Prognostic Biomarkers

To check the relevance of the PRER genes, we check whether the patients with different feature values are different in terms of their survival distribution. For a PRER protein pair, A-B, we group patients such that those patients with protein expression values A>B are in one group and the rest are in the other group. We then check if the survival distribution of these groups is different using the log-rank test. The Figure 31shows the age and sex adjusted KM plots for each cancer type and the top-ranked feature.

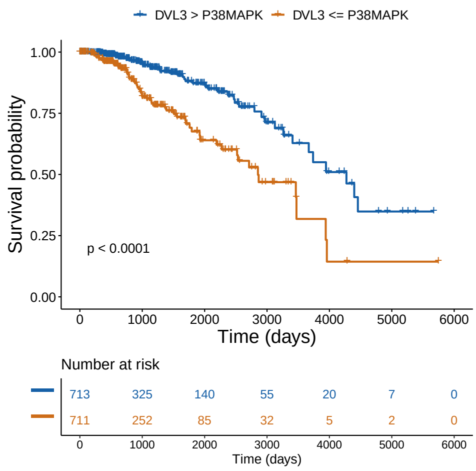


(a) BRCA

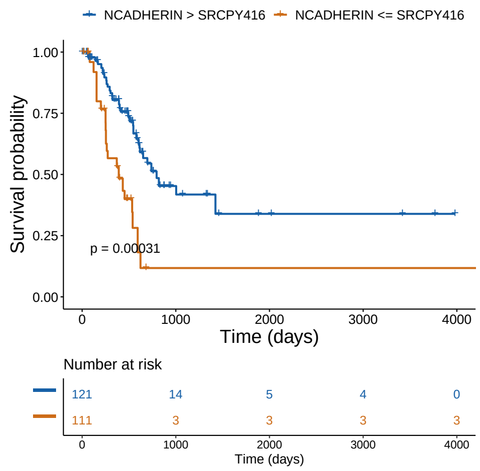


(b) BLCA

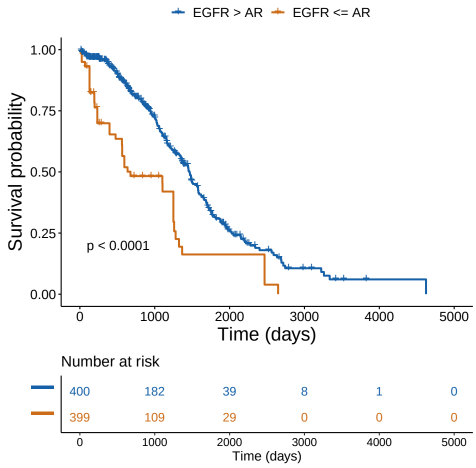


(c) OV

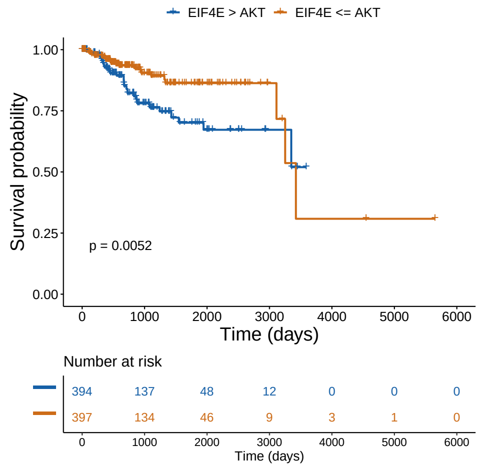


(d) UCEC

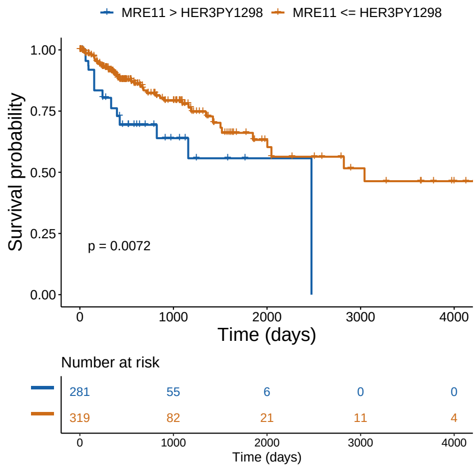


(e) COAD

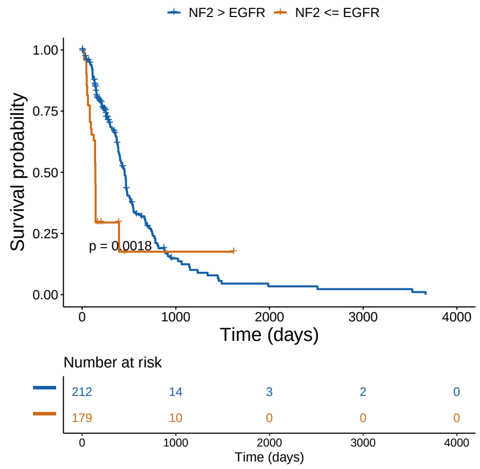


(f) GBM

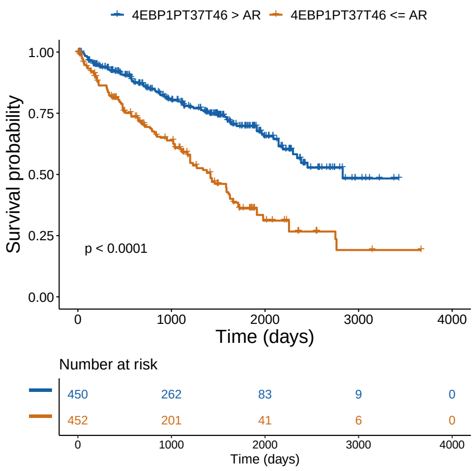


**(g)** KIRC

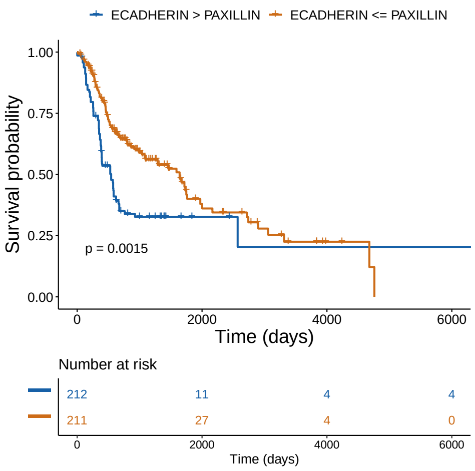


**(h)** HNSC

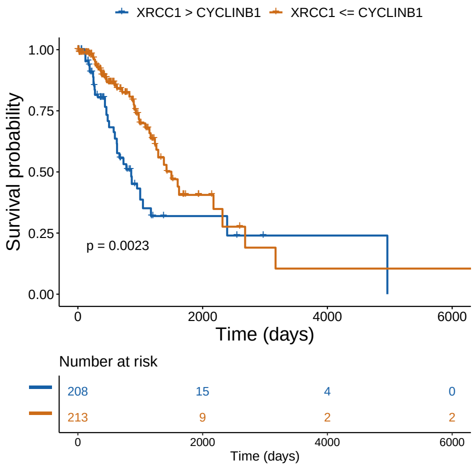


**(i)** LUAD

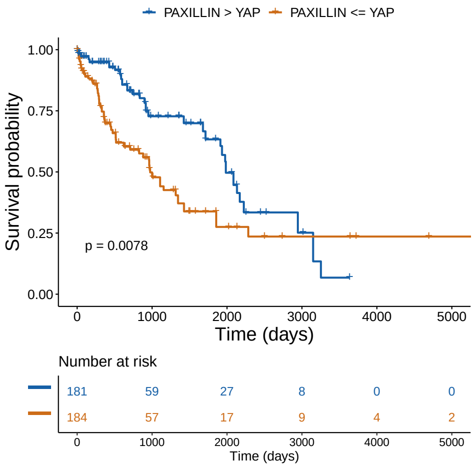


**(j)** LUSC

**Figure 31:** Age and sex adjusted Kaplan-Meier plot for each cancer type based on overall survival. Number at risk denotes the number of patients at risk at a given time, and p-value is calculated with the log-rank test.
